# SORLA upregulation suppresses global pathological effects in aged tauopathy mouse brain

**DOI:** 10.1101/2025.06.25.661599

**Authors:** Huijie Huang, Christina Huan Shi, Wenqi Yang, Juan Piña-Crespo, Jay Bhatnagar, Julian Curatolo, Rabi Murad, Palak Shah, Alex Campos, Alexandra Houser, Rebecca A. Porritt, Tongmei Zhang, Qiang Xiao, Shengjie Feng, Kevin Y. Yip, Timothy Y. Huang

## Abstract

A role for the trafficking receptor SORLA in reducing Aβ levels has been well-established, however, relatively little is known with respect to whether and how SORLA can potentially affect tau pathology *in vivo*. Here, we show that transgenic SORLA upregulation (SORLA TG) can reverse pathological effects in aged PS19 (P301S tau) mouse brain, including tau phosphorylation and seeding, ventricle dilation, synapse loss, LTP impairment and glial hyperactivation. Proteomic analysis indicates reversion of PS19 profiles in PS19/SORLA TG hippocampus, including pathological changes in synapse-related proteins as well as key drivers of synaptic dysfunction such as Apoe and C1q. snRNA-seq analysis reveals suppression of PS19- signatures with SORLA upregulation, including proinflammatory induction of *Plxnb1/Plxnb2* in glia. Tau seeding and aggregation, neuroinflammation, as well as PlxnB1/B2 induction are exacerbated in PS19 hippocampus with SORLA deletion. These results implicate a global role for SORLA in neuroprotection from tau toxicity in PS19 mouse brain.

## Introduction

Alzheimer’s disease (AD) is an age-related dementia disorder affecting the elderly. In AD, beta-amyloid (Aβ) plaques, and neurofibrillary tangles (NFTs) comprising hyperphosphorylated tau accumulate in brain. While most AD cases occur sporadically, a minority of cases (<5%) (Bekris, Yu et al. 2010) represent an inherited familial form of AD with early onset symptoms in people under 65 years old. Individuals carrying genetic variants of the amyloid precursor protein (APP), as well as genes encoding APP processing machinery within the γ- secretase complex (PS1 and PS2) manifest early-onset AD at near-complete levels of penetrance (Blacker and Tanzi 1998). Whether other causal genetic factors can influence sporadic AD remains unclear.

Recent genome wide association studies (GWAS) have linked multiple gene variants with altered AD risk, including the class I membrane receptor endosomal trafficking factor, SORLA, or “Sortilin-related receptor containing LDLR class A repeats” (*SORL1*, LR11) (Lambert, Ibrahim-Verbaas et al. 2013, Kunkle, Grenier-Boley et al. 2019). SORLA is a type I transmembrane trafficking component comprising N-terminal VPS10, EGF-like/YWTD, LDLR and FN3 repeats within the ectodomain, as well as a short cytoplasmic tail (Andersen, Rudolph et al. 2016). SORL1 variants were initially identified in SORL1 locus in association with late-onset AD, and siRNA-mediated SORLA downregulation enhanced amyloidogenic APP trafficking and processing (Rogaeva, Meng et al. 2007). Subsequent studies have further demonstrated the association of SORL1 coding variants with late onset AD (Vardarajan, Zhang et al. 2015, Cuccaro, Carney et al. 2016, Holstege, van der Lee et al. 2017, Louwersheimer, Cohn-Hokke et al. 2017). Interestingly, SORLA variants have also been identified in inherited late-onset AD (Fernandez, Black et al. 2016), and familial SORLA variants have also been linked to early onset AD in European populations (Pottier, Hannequin et al. 2012, Nicolas, Charbonnier et al. 2016, Verheijen, Van den Bossche et al. 2016). Large-scale exome-sequencing studies implicate a significant correlation between loss-of-function SORLA variants and AD onset (Raghavan, Brickman et al. 2018), and growing evidence suggests that some SORLA variants such as the dimerization-deficient Y1816C (Jensen, Raska et al. 2024) or R953C (Fazeli, Child et al. 2024) may be linked to dominant causality in AD (Scheltens, De Strooper et al. 2021).

Reduced SORLA expression was originally linked to AD (Scherzer, Offe et al. 2004), while deletion of murine SORLA (*Sorl1)* showed enhanced Aβ accumulation in cortex (Andersen, Reiche et al. 2005). In contrast, transgenic expression of human SORLA reduces Aβ accumulation and pathology in APP/PS1 AD mice (Caglayan, Takagi-Niidome et al. 2014). SORLA is a component of the retromer endosomal trafficking complex and has been observed to bind VSP26 (Fjorback, Seaman et al. 2012) and SNX27 (Huang, Zhao et al. 2016) subunits of the retromer complex within the C-terminal tail domain. Expression of retromer components have been shown to be downregulated in AD (Small, Kent et al. 2005). VPS26 haploinsufficiency in mice have been shown to elevate amyloidogenic APP processing, Aβ accumulation, and induce synaptic defects (Muhammad, Flores et al. 2008), and pharmacological stabilization of the retromer complex can attenuate amyloidogenic APP processing in primary cultured mouse neurons (Mecozzi, Berman et al. 2014). Interestingly, deletion of a neuron-specific VPS26 isoform (VPS26b) in mouse brain is associated with decreased total and cell-surface SORLA levels, as well as increased murine brain Aβ and CSF tau (Simoes, Guo et al. 2021), implicating interdependent roles for retromer, SORLA and AD pathology.

SORLA-mediated neuroprotective effects involve reduction in Aβ levels through various mechanisms, including trafficking of APP from the endosome to the Golgi (Fjorback, Seaman et al. 2012) or cell surface (Huang, Zhao et al. 2016), thus attenuating amyloidogenic APP cleavage by BACE1 at late endosomes. In addition, SORLA reduces APP and BACE1 (Spoelgen, von Arnim et al. 2006) interactions while directly binding and facilitating Aβ internalization and lysosomal degradation (Caglayan, Takagi-Niidome et al. 2014, Liu, Zhu et al. 2020). Neuroprotective effects SORLA on Aβ accumulation and generation are well documented, however, its effects on tau pathology and proteotoxicity still remains somewhat unclear.

Here, we provide evidence that SORLA upregulation prevents pathological ventricular enlargement, tau phosphorylation and seeding, synaptic loss, impaired synaptic plasticity and glial hyperactivation in transgenic PS19 mouse hippocampus expressing human P301S tau. Proteomic analysis revealed shifts in synaptic proteins and microglial activation state in tau-burdened hippocampus compared to wildtype (WT) animals; pathological changes in proteins including key drivers of tau pathology such as Apoe and C1q were reduced with SORLA upregulation. Further, induction of disease-associated microglia (DAM) (Keren-Shaul, Spinrad et al. 2017), and astroglia (DAA) (Habib, McCabe et al. 2020) signatures in tau-burdened hippocampus were attenuated by SORLA upregulation by snRNA-seq analysis. snRNA-seq changes in transcripts related to the synapse, cell-cell communication and signal transduction in PS19 oligodendrocytes and neurons were also reversed with SORLA upregulation. SORLA upregulation could reverse reduced expression of synaptic components such as Nrxn1 in astrocytes, neurons and microglia of PS19 hippocampus, and while SORLA upregulation enhanced uptake of tau oligomers in cultured neurons and microglia, enhancement of tau trafficking to late/endosomes and lysosomes were observed in microglia. In addition, SORLA suppressed pathological upregulation of Sema4D receptors, PlxnB1 and PlxnB2, while its deletion increased PlxnB1/PlxnB2 levels in PS19 hippocampus. Together, these results demonstrate that SORLA upregulation reverses pathological changes in synaptic structure, function and plasticity in tau-burdened hippocampus and implicates enhanced SORLA expression as an upstream driver for global neuroprotective effects in a tauopathy mouse model.

## Results

### SORLA upregulation attenuates pathological features in aged PS19 hippocampus

The PS19 (P301S tau) mouse, a widely used model of tau pathology, manifests pathological features by 8-9 months of age (Figure 1A) (Yoshiyama, Higuchi et al. 2007). We crossed PS19 animals with a mouse line expressing human SORLA under the regulation of a CAG promoter at the Rosa26 locus (SORLA TG) (Caglayan, Takagi-Niidome et al. 2014). By comparing AT8-reactive (pS202/T205) ptau levels in soluble hippocampal lysates in 3- and 9-month-old (MO) animals, we observed low AT8 ptau levels in 3MO PS19 hippocampus and SORLA TG/PS19 animals (Figure 1B, C). At 9MO, however, AT8 ptau levels were significantly elevated in PS19 animals compared to WT, and significantly reduced in SORLA TG/PS19 hippocampus compared to PS19 (Figure 1B,C). At 3MO, we observed little or no change in GFAP levels, an astrogliosis marker, in P301S tau animals; however, GFAP levels were significantly higher in 9MO PS19 hippocampus and reduced in SORLA TG/PS19 animals (Figure 1B, C). Increased GFAP levels tightly correlated with AT8 ptau induction in 9MO PS19 hippocampus (Figure 1B); animals with increased AT8 ptau levels also showed proportionally elevated GFAP levels (Figure 1D). We observed a high degree of variability in AT8 levels in PS19 hippocampus (Figure 1C) similar to other studies characterizing PS19 (DeVos, Miller et al. 2017, Soni, Lin et al. 2024) or PS19 mice combined with human APOE4 knock-in alleles (Shi, Manis et al. 2019, Koutsodendris, Blumenfeld et al. 2023, Seo, O’Donnell et al. 2023, Carling, Fan et al. 2024). We next compared effects of SORLA upregulation on insoluble tau and AT8 ptau levels in 11MO PS19 and SORLA TG/PS19 (TG_PS19) cortex. Similar to our observations in hippocampus (Figure 1B, C), we observed significant reduction of soluble AT8 ptau tau levels normalized over total tau in SORLA TG/PS19 compared to PS19 cortex (Figure 1E), however, trending decreases in insoluble tau and AT8 ptau and AT8 ptau/tau levels were observed in SORLA TG/PS19 over PS19 animals (Figure 1E). We also determined whether SORLA upregulation could alter seed-competent tau in 11MO cortex samples using an HEK293 tau-RD biosensor assay system (Holmes, Furman et al. 2014). Using soluble lysates from PS19 and SORLA TG/PS19 lysates (Figure 1F, G), we observed reduced tau seeding FRET activity in SORLA TG/PS19 cortical lysates in HEK tau-RD cells (Figure 1H). These results together indicate that SORLA upregulation can reduce AT8 ptau and seed-competent tau in PS19 brain.

**Figure 1.**
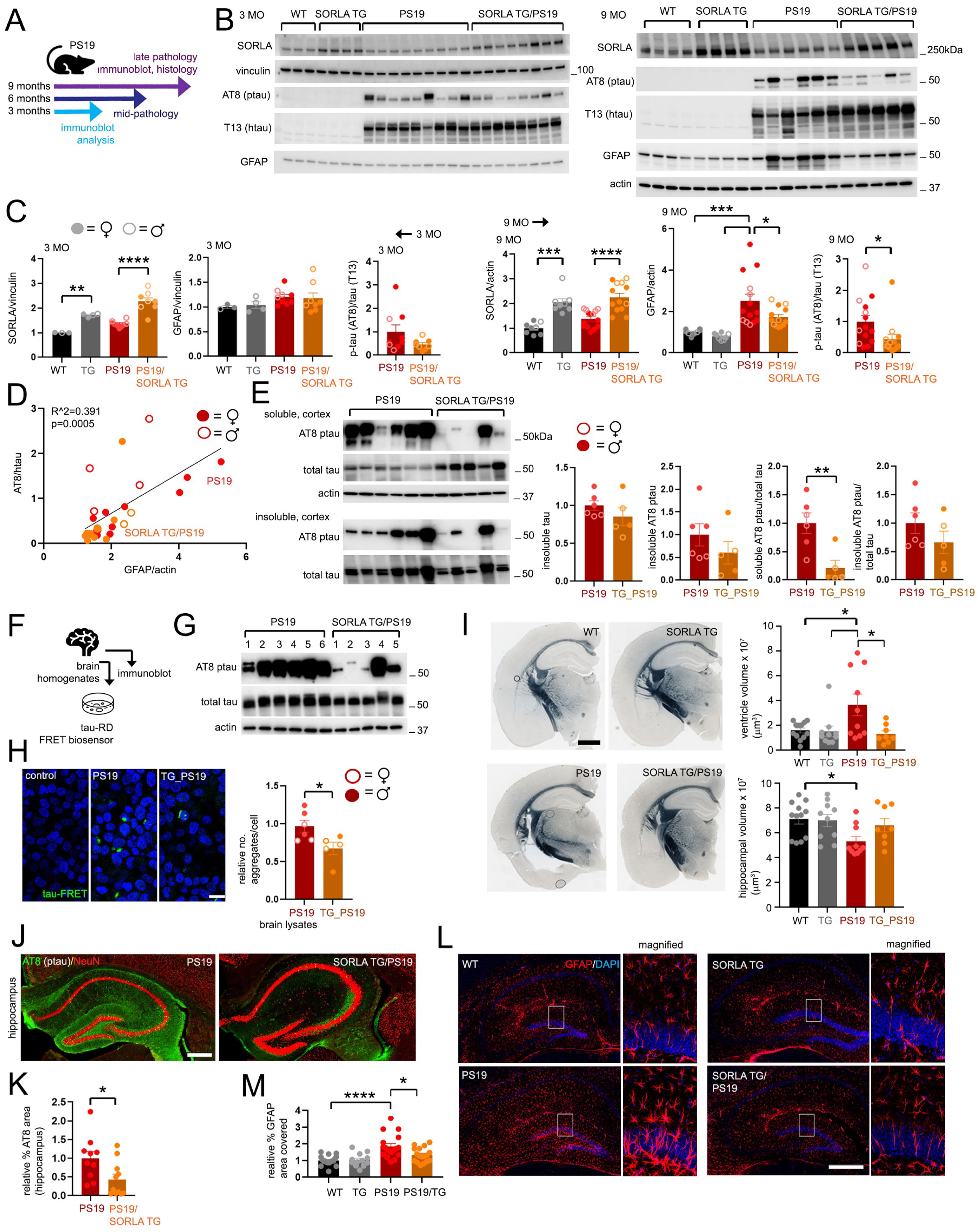
SORLA upregulation attenuates pathological changes in PS19 hippocampus. (A) Schematic depicting experimental workflow and pathological stages in PS19 mouse brain. (B) Immunoblot analysis of soluble hippocampal lysates from 3 month-old (MO) (left blots) and 9 MO wildtype (WT), SORLA TG, PS19 and SORLA TG/PS19 animals (right blots). Lysates were immunoblotted for SORLA, vinculin/actin, pS203/T205 tau (AT8), total human tau (T13) or GFAP as indicated. (C) Graphs depicting quantified band intensities from immunoblots in (B). Individual plots represent individual animals at 3 MO (left graphs) and 9 MO (right graphs). Relative band intensity values were normalized to WT (SORLA, GFAP) or PS19 (AT8/T13 tau) animals (WT or PS19 levels were set to 1.0). (D) Correlation between AT8 ptau and GFAP levels in 9 MO hippocampus from relative band intensity values quantified in (C). (E) AT8 ptau and total tau from soluble and insoluble fractions in PS19 and SORLA TG/PS19 (TG_PS19) 11MO cortex. Adjacent graphs depict insoluble tau or AT8 ptau (left graphs) or soluble/insoluble AT8 ptau/total tau ratios (right graphs) normalized to PS19 (set to 1.0). (F-H) Characterizing seed competent tau in PS19 and SORLA TG/PS19 brain. Experimental scheme (F): soluble 11MO PS19 and SORLA TG/PS19 cortical brain extracts were subjected to immunoblot analysis (G) or applied to HEK293 tau-RD cells (H) and assayed for tau-FRET activity. (H) Representative tau-FRET images (green) or DAPI (blue) are shown, bar=20um. Adjacent graph depicts relative number of tau-FRET aggregates from PS19 or TG_PS19 animals, normalized to PS19 (set to 1.0); extracts derived from female or male animals are indicated. (I) Sudan black stained coronal sections from 9 MO WT, SORLA TG, PS19 and SORLA TG/PS19 animals, bar = 1mm. Ventricle dilation was quantified in coronal sections, ventricle volume (upper graph, um^3^ x 10^7^) and hippocampal volume (lower graph) in 30um slices is shown for individual animals (WT: 7F, 5M; STG: 7F, 3M; PS19: 7F, 3M; STG/PS19: 5F, 3M). (J) Representative images from histological hippocampal sections from 9 MO PS19 or SORLA G/PS19 animals stained for AT8 ptau (green) and NeuN (red), bar = 1mm. (K) Quantification of AT8 ptau area from stained sections in (J), relative stained area was normalized to PS19 animals (set to 1.0). (PS19: 5F, 5M; STG/PS19: 7F, 4M). (L) Representative images from 9 MO hippocampal sections stained for GFAP (red) or nuclei (DAPI, blue), bar = 500um. (M) Quantification of percentage GFAP staining area from images in (L). (WT: 7F, 7M; STG: 6F, 6M; PS19: 10F, 7M; STG/PS19: 10F, 7M). All graphs represent mean ± SE. Statistical significance was determined by One-way ANOVA in (C) (SORLA, GFAP comparisons), (I) and (M), or unpaired Student’s t-test in (C) (AT8/T13 comparisons), (E), (H) and (K), *p<0.05, **p<0.01, ***p<0.001, ****p<0.0001. Empty and filled plots represent female and male animals in (C), (D), (E) and (H) as indicated.

Next, we examined effects of SORLA upregulation on histopathological changes in P301S tau brain. As reported by others (Yoshiyama, Higuchi et al. 2007), we observed ventricle enlargement in coronal sections in 9MO PS19 animals, whereas SORLA upregulation in PS19 animals largely suppressed ventricle dilation (Figure 1I). Moreover, SORLA upregulation suppressed accumulation of AT8 ptau in PS19 hippocampus (Figure 1J, K), as well as indicators of astrogliosis including increased GFAP staining and astrocyte hypertrophy (Figure 1L, M). Together, these results indicate that SORLA upregulation can suppress pathological changes in aged PS19 mouse brain.

### SORLA upregulation suppresses proteomic changes in PS19 hippocampus

We next determined whether SORLA upregulation in SORLA TG animals could affect bulk proteomic alterations in PS19 mouse hippocampus. To this end, we performed label-free mass spectrometry analysis of protein lysates from 9MO WT, SORLA TG (“TG”), PS19 and SORLA TG/PS19 (“TG_PS19”) mouse hippocampus. Principal Component Analysis (PCA) revealed good clustering of replicate animals from each genotype (Figure 2A), with the highest number of differentially expressed proteins (DEPs) identified when comparing PS19 vs WT hippocampus (Figure 2B, C, and Figure S1A, B; Table S1). We also observed proteins specifically expressed in PS19 hippocampus; 227 DEPs were PS19-specific (altered specifically in PS19 vs WT, not in TG_PS19 vs WT animals) (Figure 2D, E), and 140 PS19-specific DEPs were observed in PS19 vs TG comparisons (Figure S1C, D).

**Figure 2.**
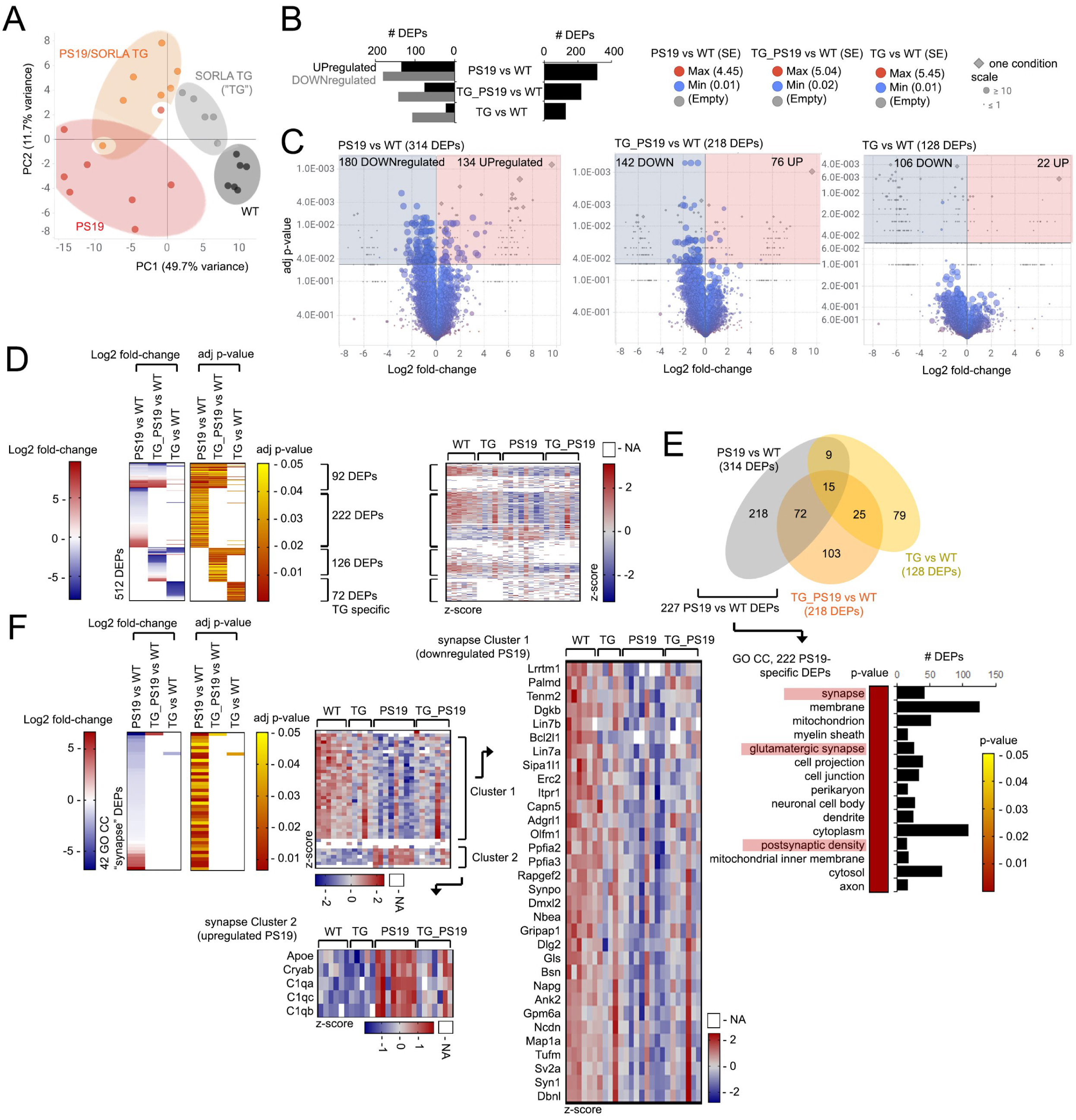
Proteomic changes in PS19 hippocampus are reversed with SORLA upregulation. (A-C) Label-free proteomic analysis of 9MO mouse hippocampus. Protein lysates from WT (black; n=6, 4F, 2M), SORLA TG (“TG”, gray; n=5, 3F, 2M), PS19 (red; n=8, 5F, 3M), and SORLA TG/PS19 (“TG_PS19”, orange; n=7, 4F, 3M) animals were subjected to label-free proteomic analysis, and DEPs (adjp<0.05) were subjected to PCA analysis in (A). PCA clusters are highlighted in colored ovals (A). (B) Total number (right graph) and number of upregulated (black bars) and downregulated (gray bars) DEPs (left graph), and (C) volcano plots depicting Log2 fold-change and adj p-values in “PS19 vs WT”, “TG_PS19 vs WT”, and “TG vs WT” hippocampus. Number of DEPs (shown in brackets), as well as standard error (SE), frequency of detection are indicated by scaled circles; proteins absent under one condition are indicated by gray diamonds (legend) in (C). (D) Log2 fold-change and adj p-value in DEPs in comparisons indicated (left heatmaps), and z-score distribution for individual animals of the genotypes indicated (right heatmaps); “NA” values from missing peptide values are indicated in z-score heatmaps in white. (E) Venn diagram depicting overlap between “PS19 vs WT”, “TG_PS19 vs WT” and “TG vs WT” DEPs. 227 DEPs detected in PS19 vs WT (and absent in TG_PS19 vs WT) were subjected to GO analysis, p-value (heatmap) and number of DEPs of top 15 GO CC categories (categories related to synaptic and neuronal structures highlighted) shown. DEPs with opposing up or downregulation trends will count as unique DEPs in more than one comparison group, and so the total DEPs may not match the sum of all Venn subgroups. (F) Log2 fold-change and adj p-value (left heatmaps) and z-score (right heatmaps) distribution of 42 GO CC “synapse”- related DEPs specific to PS19 vs WT comparisons. Z-score distribution of GO CC “synapse” DEPs from (E) that feature downregulation (Cluster 1) or upregulation in PS19 (Cluster 2) are indicated. See also Figure S1.

Gene Ontology (GO) analysis of the 227 PS19-specific DEPs revealed enrichment in GO Cellular Component (CC) categories related to synapses, with GO CC “synapse” (GO:0045202) as the top CC category enriched in PS19-specific DEPs (Figure 2E, Table S2). Of the 42 GO CC “synapse”-related DEPs, DEPs were downregulated in PS19 compared to other genotypes in one cluster (Cluster 1, Figure 2F), whereas one cluster featured DEPs induced in PS19 hippocampus compared to other groups (Cluster 2, Figure 2F). Numerous GO CC “synapse” categories were also identified PS19 vs TG hippocampus (Figure S1F, G). In comparing proteomic profiles in PS19 vs TG and TG_P19 vs TG groups, we observed enrichment of C1qa, C1qb and C1qc in the GO KEGG “complement and coagulation cascades” pathway; Apoe, Cryab, and C1qa, C1qb and C1qc were also enriched in GO CC PS19-specific (PS19 vs TG) DEPs (Figure S1F, G; Table S2). Interestingly, we observed robust upregulation of synapse-related DEPs such as Apoe, Cryab and C1qa, C1qb and C1qc in PS19 which was normalized to levels comparable to WT or TG hippocampus in TG_PS19 animals (Figure 2F and Figure S1E, G). We also performed GO analysis on PS19 vs WT or TG_PS19 vs WT DEPs and searched for enrichment in synaptic DEPs using SYNGO (syngoportal.org) (Koopmans, van Nierop et al. 2019). PS19 vs WT featured more synaptic DEPs compared to TG_PS19 vs WT, indicating that SORLA upregulation reverses changes in synaptic protein levels observed in PS19 hippocampus (Figure S2A). Numerous GO Biological Process (BP) categories were identified and enriched in PS19 vs WT hippocampus related to the structure and function of synapses, including presynaptic and postsynaptic elements (Figure S2B, C). These results suggest that genes related to synaptic maintenance and function were potentially altered in PS19 hippocampus, and largely attenuated in SORLA TG/PS19 animals.

### SORLA upregulation attenuates microglial activation and drivers of pathological and synaptic dysfunction in PS19 hippocampus

Alterations in microglia activation state have been observed in PS19 mouse brain, and various DAM signatures (Keren-Shaul, Spinrad et al. 2017) have been shown to be induced in PS19 microglia (Wang, Fan et al. 2022). We observed induction of DAM-like DEPs in PS19 hippocampus, including Fth1, Ctsd, Apoe, Ctsz and Cd63, that were largely attenuated in SORLA TG/PS19 animals (Figure 3A). As this suggests SORLA upregulation suppresses microglia activation in PS19 brain, we examined whether markers of microglia activation such as Cd68 were similarly reduced in SORLA TG/PS19 hippocampus. Indeed, Cd68, Iba1 and Clec7a (Figure 3B-E and Figure S2D, E) staining in 9MO SORLA TG/PS19 hippocampus was significantly reduced compared to PS19. In addition, we observed downregulation of the microglia homeostatic marker P2ry12 in PS19 hippocampus which was normalized in SORLA TG/PS19 animals (Figure S2F, G). Thus, SORLA upregulation can suppress microglia activation associated with tau proteotoxicity in PS19 hippocampus. Whether this effect is due to SORLA modulation in microglia, tau proteotoxicity in neurons or a combination of both features remains unclear.

**Figure 3.**
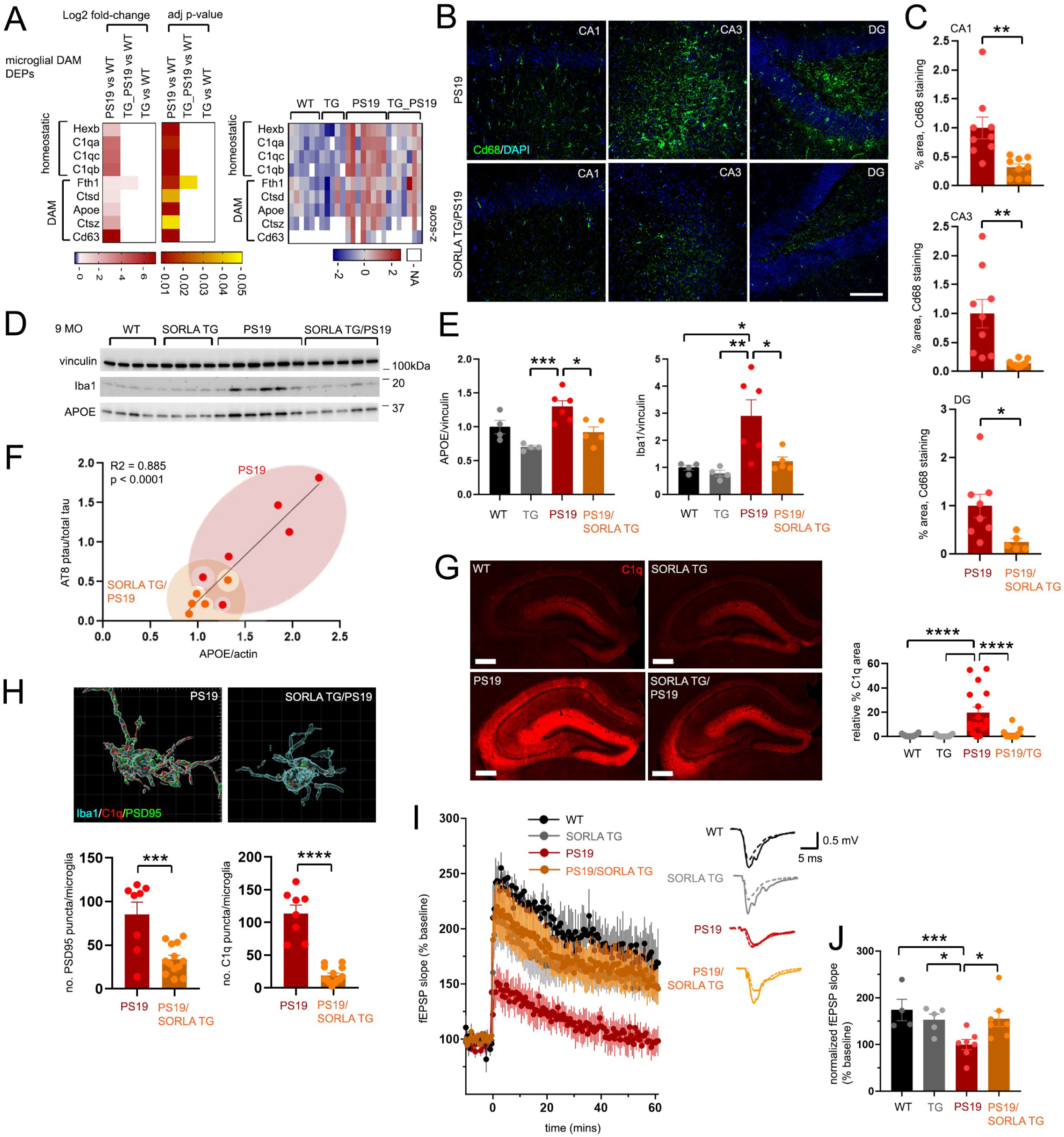
SORLA upregulation attenuates microglia activation and synaptic impairment in PS19 hippocampus. (A) Log2 fold-change/adj p-value and z-score distribution of DAM-related DEPs identified in 9 MO mouse hippocampus by label-free proteomic analysis. (B) Representative images of hippocampal sections from 9 MO PS19 or SORLA TG/PS19 animals stained to visualize Cd68 (green) or nuclei (DAPI, blue), bar = 100um. Images were obtained from DG, CA3 and CA1 regions as indicated. (C) Graphs depicting quantified percentage Cd68 staining area from PS19 and SORLA TG/PS19 animals is shown from imaged regions in (B). (PS19: 6F, 3M; STG/PS19: 6F, 3M for CA1, CA3). (D) Immunoblot analysis of hippocampal lysates from 9MO WT, SORLA TG, PS19 and SORLA TG/PS19 animals. Lysates were immunoblotted for vinculin, Iba1, and Apoe as indicated. (E) Graphs depicting quantified band intensities of immunoblots shown in (D), normalized to WT (set to 1.0). (WT: 4F; STG: 4F; PS19: 6F; STG/PS19: 5F) (F) Correlation between APOE expression and AT8 ptau expression in 9MO hippocampus. Plots for PS19 (red) and SORLA TG/PS19 (orange) are shown, only females were matched for correlation analysis. (G) Representative hippocampal sections from 9MO WT, SORLA TG, PS19 and SORLA TG/PS19 animals stained for C1q (red), bar = 400um. (H) Graph depicts quantified C1q fluorescence intensity normalized to WT (set to 1.0). (WT: 8F, 6M; STG: 8F, 4M; PS19: 10F, 7M; STG/PS19: 10F, 7M). (I) Representative reconstructed 3D confocal images of C1q (red) and PSD95 (green) puncta localized within microglia (blue) in PS19 and SORLA TG/PS19 hippocampus. (J) Graphs depict quantification of PSD95 (left graph) and C1q (puncta) internalized per microglia in PS19 or SORLA TG/PS19 hippocampus. (PS19: 5F, 3M; STG/PS19: 9F, 4M). (K) SORLA upregulation rescues LTP impairment in PS19 hippocampus. Acute slices from WT (black), SORLA TG (gray), PS19 (red) or SORLA TG/PS19 (orange) 6.5 to 7.5 MO animals were subjected to LTP induction (time 0) in the presence of 100 µM picrotoxin and mean fEPSP slopes over baseline was quantified from recordings in the dentate gyrus (DG) region. Baseline recordings were taken 20 mins. before and 60 mins. after stimulation. Adjacent fEPSP trace recordings (mean±SE, averaged for each animal) before (solid lines) and after (dotted lines) LTP induction for each experimental genotype. (L) Cumulative fEPSP slopes during the last 10 mins. of recording (50-60 mins. after induction) were averaged from from individual animals of each genotype (WT: 2F, 2M, n=4mice /6 slices; SORLA TG: 3F, 2M, n=5 mice/6 slices; PS19: 3F, 4M, n=7 mice/12 slices, SORLA TG/PS19: 5F, 2M, n=7 mice/13 slices). (M, N) Effects of SORLA modulation on tau uptake in microglia. (M) Representative images of WT and SORLA TG (TG) microglia following 2h tau oligomer (15 nM) treatment; microglia were fixed and stained for endolysosomal markers (EEA1, Rab7, LAMP1; green), T13 tau (red) or nuclei (DAPI, blue), bar=10um. (N) Quantification of tau uptake in WT and TG microglia (upper graph). Lower graphs depict EEA1, Rab7 and LAMP1 co-localization with tau following exposure to tau oligomers using Pearson’s correlation values. Plots represent values from one imaged field, and all values were derived from three independent experiments. All graphs depict mean ± SE. Statistical significance was determined by unpaired Student’s t-test (C), (J), (N) or One-way ANOVA (E), (H), (L), or simple linear regression (F), *p<0.05, **p<0.01, ***p<0.001, ****p<0.0001. See also Figure S2.

Apoe and C1q are key drivers of pathological degeneration in PS19 brain, with previous studies showing deletion of Apoe (Shi, Manis et al. 2019) or C1q (Dejanovic, Wu et al. 2022) can suppress degenerative features such as ventricular dilation PS19 mouse brain. Since Apoe and C1q induction in PS19 hippocampus was suppressed with SORLA upregulation (Figure 3A), we validated effects of SORLA upregulation on Apoe and C1q by immunoblot and histological analysis. We found Apoe (Figure 3D-F) and C1q (Figure 3G, H) were induced in 9MO hippocampus, and consequently normalized to near-WT levels in SORLA TG/PS19 animals (Figure 3D-H). Effects of SORLA upregulation on Apoe appeared to be PS19 dependent; Apoe levels did not appear to differ significantly between SORLA TG and WT hippocampus (Figure 3A, D and E). We also observed tight correlation between AT8 ptau levels and APOE in PS19 and SORLA TG/PS19 hippocampus (Figure 3F). Both Apoe and C1q are also biologically associated with synaptic function; C1q is a component of the complement pathway that accumulates at synapses to promote selective synaptic pruning during brain development (Huttenlocher 1979, Feinberg 1982) and mediates pathological elimination of synapses where microglia engulf or “prune” synapses (Hong, Beja-Glasser et al. 2016). APOE has been implicated in synaptic removal by colocalizing and enhancing Aβ accumulation at synapses (Koffie, Hashimoto et al. 2012), and APOE4 depletion in astrocytes suppresses microglial synaptic removal in an APOE4 KI model of mouse tauopathy (Wang, Xiong et al. 2021).

Given that SORLA upregulation can normalize expression of synaptic components in PS19 brain we examined whether pre- and post-synaptic junctions were altered in PS19 and SORLA TG/PS19 hippocampus. In agreement with previous reports in PS19 animals (Yoshiyama, Higuchi et al. 2007, Dejanovic, Huntley et al. 2018), we observed decreased colocalization of PSD95 (postsynaptic)/Vglut1 (presynaptic) puncta in 9MO PS19 hippocampus compared to WT or SORLA TG, where SORLA TG/PS19 hippocampus featured partial restoration of colocalizing PSD95/Vglut1 particles (Figure S2H, I). Since SORLA upregulation can reverse pathological microglia activation markers in PS19 brain, we also explored whether microglial PSD95 and/or C1q uptake was altered with SORLA transgene expression (Figure 3I, J). Internalized C1q and PSD95 puncta in PS19 microglia was largely attenuated in SORLA TG/PS19 animals, further indicating that SORLA upregulation can reverse pathological C1q and PSD95 uptake in microglia (Figure 3I, J). When comparing long-term synaptic plasticity (LTP) in acute hippocampal slices from WT, SORLA TG, PS19 and SORLA TG/PS19 animals, we observed that impaired LTP response in PS19 DG was largely restored in SORLA TG/PS19 animals (Figure 3K, L), demonstrating that SORLA upregulation can largely restore functional synaptic impairment in PS19 hippocampus.

In addition to alternations in C1q/PSD95 uptake, we also explored whether SORLA upregulation could affect tau uptake and trafficking in microglia. We found that 2h following exposure to tau oligomers, SORLA TG microglia exhibited enhanced tau uptake compared to WT (Figure 3M, N). We also observed a trending enhancement of tau colocalization with early endosomes (EEA1), and significantly enhanced tau colocalization with late endosomes and lysosomes in SORLA TG microglia (Figure 3M, N). This suggests that while SORLA upregulation can attenuate synaptic uptake *in vivo*, SORLA upregulation can also enhance uptake of tau oligomers *in vitro*.

### SORLA upregulation reverses cell-specific pathological transcriptomic glial profiles in PS19 hippocampus

Our results indicate SORLA upregulation can reverse pathological effects in aged PS19 brain. To explore whether these restorative effects correlate with cell-specific transcriptomic changes, we subjected hippocampal tissue from 9MO WT, PS19 and SORLA TG/PS19 (“TG_PS19”) to single nucleus RNAseq (snRNA-seq) analysis. We sequenced a total of 32,048 single nuclei, clustered into 18 main clusters of varying cell identity (Figure 4A, B and Figure S3A-B), retaining 28,348 high quality single nuclei data after doublet removal and quality control verification. We determined cell cluster identities using expression of cell type markers, as well as reference mapping to the Allen Brain Cell Atlas. We then reclustered nuclei from microglia (Cluster 3), astrocytes (Clusters 1,14), neurons (Clusters 2,6,10,11,15) and compared expression profiles between WT, PS19 and TG_PS19 groups. As expected, we did not detect human SORLA (SORL1) expression in WT and PS19 samples, and we observed upregulation of human SORLA (SORL1) in neurons, glial cell types such as astrocytes, oligodendrocytes, oligodendrocyte precursor cells (OPC’s) and microglia, as well as other cell types including pericytes, VLMC and endothelial cells (Figure S3C).

**Figure 4.**
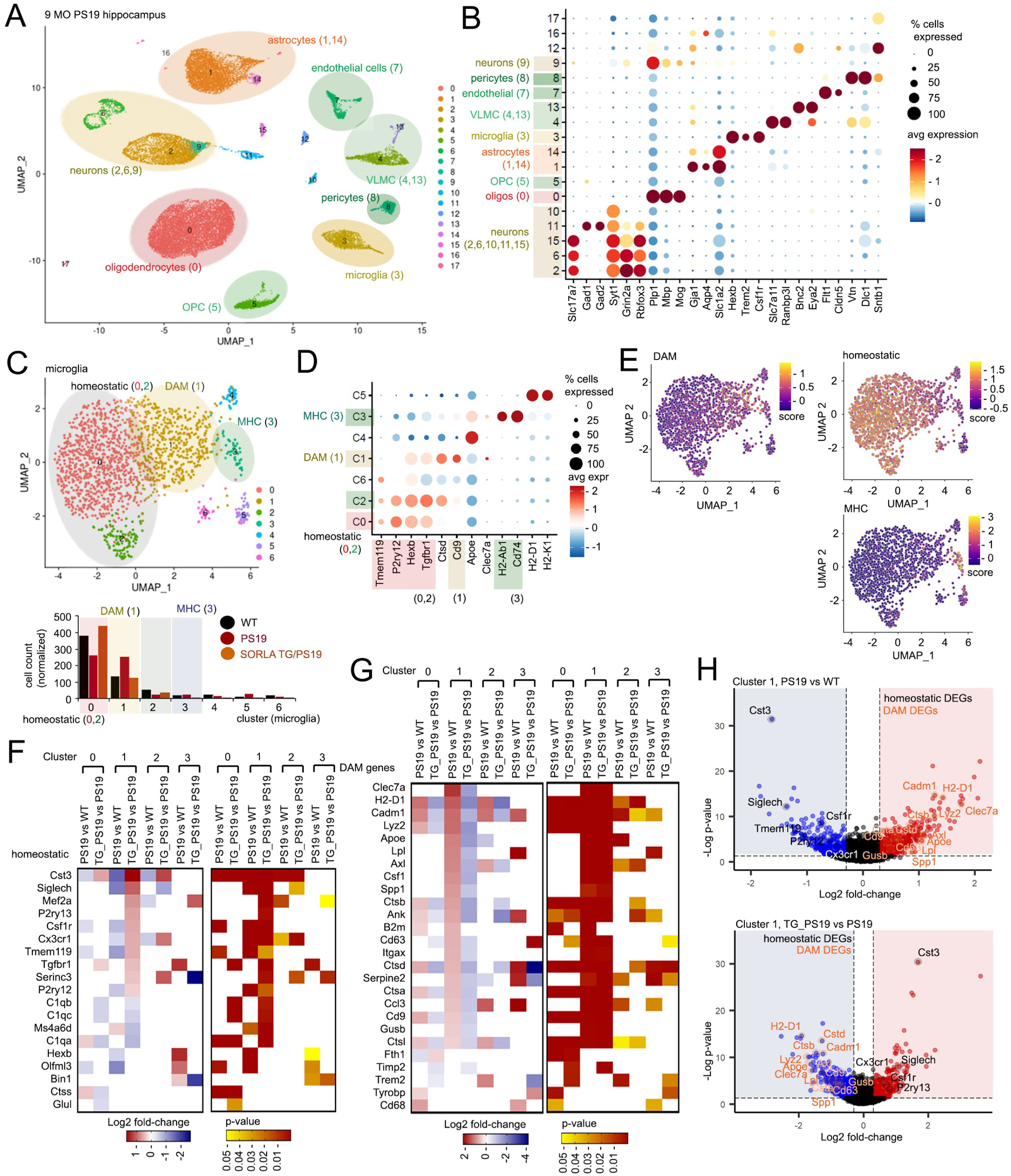
**snRNAseq analysis SORLA upregulation suppresses pathological induction of microgial DAM signatures in PS19 hippocampus**. (A) Umap distribution of 29,438 nuclei clustered from 9MO WT, PS19 and SORLA TG/PS19 hippocampus. Cell types identified according to alignment with murine reference profiles from the Allen Brain Atlas are indicated. (B) Dot plot depicting expression profiles of key identifying trancriptomic signatures from each cluster from (A), heat scales for expression levels, as well as % expression (dot size) are shown. (C) Umap distribution of microglia nuclei reclustered from microglia Cluster 3 in (A). Homeostatic (0, 2), DAM (1) and MHC (3) clusters are indicated. Lower graph depicts normalized cell numbers in WT, PS19 and SORLA TG/PS19 microglia in the clusters indicated. (D) Dot plot depicting expression of microglia gene signatures in microglia clusters 0 through 6. (E) Heat-scaled Umap depicting expression of homeostatic, DAM and MHC genes within reclustered microglia; color scale represents enrichment of genes within each category using scores generated. (F, G) Microglia homeostatic/DAM DEGs and their expression profiles. Log2 fold-change (left heatmaps) and p-value (right heatmaps) of DEGs associated with homeostatic (F) and DAM (G) microglia in PS19 vs WT and TG_PS19 vs PS19 comparisons are shown. (H) Volcano plots depicting up and downregulated DEGs in PS19 vs WT and TG_PS19 vs PS19 microglia comparisons in microglia Cluster 1 (p-value<0.05; Log2 fold-change<-0.3, >0.3). Homeostatic (black) and DAM (orange) DEGs featuring opposing expression profiles in PS19 and TG_PS19 microglia are indicated. See also Figure S3.

Transcriptomic profiles from reclustered microglia nuclei segregated into 7 clusters (Figure 4C), with distinct separation of homeostatic (Clusters 0,2), DAM (Cluster 1) and MHC (Cluster 3) clusters (Figure 4D, E and S3D, E). Interestingly, homeostatic DEGs that were largely downregulated in PS19 DEGs (PS19 vs WT) such as *Cst3, Siglech, Mef2a, Csf1r, Cx3cr1, Tmem119*, and *P2ry12* were upregulated in SORLA TG/PS19 animals (TG_PS19 vs PS19) (Figure 4F and S3E). Trends with homeostatic DEGs were most evident in the DAM Cluster 1, while DAM-associated genes such as *Clec7a, H2-D1, Cadm1, Lyz2, Apoe, Lpl, Axl*, and *Csf1* upregulated in PS19 microglia were downregulated in SORLA TG/PS19 animals most abundantly in cluster 1 (Figure 4F, G). In agreement with our proteomics analysis, we also observed upregulation of *C1qa* in PS19 microglia which was reduced in SORLA TG/PS19 hippocampus (Figure 4F). We also observed reversed homeostatic and DAM expression gene trends in PS19 and SORLA TG/PS19 microglia in homeostatic Clusters 0 and 2 (Figure 4F-H), and upregulation of MHC-associated genes in PS19 microglia were also reversed in SORLA TG/PS19 microglia in Clusters 0, 1 and 2 3 (Figure S3F).

In addition to DEGs associated with homeostatic, DAM and MHC signatures, we observed that SORLA upregulation can reverse differentially expressed genes (DEGs) in Clusters 0 through 3 (Figure S3G). GO analysis of pathological PS19 DEGs restored in SORLA TG/PS19 microglia include DEGs associated with KEGG pathways related to “Lysosome” and “Glutamatergic synapse” (Figure S3H). KEGG lysosome-associated PS19 microglia DEGs were largely upregulated and consequently downregulated in SORLA TG/PS19 microglia (Figure S3I), indicating that SORLA upregulation can potentially normalize pathological elevation of lysosome components in PS19 microglia. SORLA upregulation also appeared to normalize numerous KEGG pathway DEGs associated with glutamatergic synapses in PS19 microglia (Figure S1J).

Reclustered astrocyte snRNA-seq profiles revealed 5 clusters (Clusters 0 through 4) (Figure 5A, B and Figure S4A, B), with disease-associated astrocytes (DAA) residing primarily in Cluster 2 (Figure 5B-D and S4C). PS19 astrocytes featured increased numbers of DAA Cluster 2 astrocytes, which were reduced in TG_PS19 hippocampus (Figure 5A, lower graph). Although expression of DAA genes were primarily enriched in Cluster 2 (Figure 5C, D), upregulated DAA signatures in PS19 astrocytes were largely reversed in TG_PS19 hippocampus in Clusters 0 through 3 (Figure 5D-F and Figure S4D). Similar to our observations in microglia, SORLA upregulation largely reversed pathological changes in PS19 astrocytes (Figure S4E); GO analysis of PS19 DEGs reversed in SORLA TG/PS19 astrocytes revealed changes in GO BP transcription (Figure S4F, G) and GO CC categories synapse categories (Figure S2F, H). We also characterized bulk changes from the main oligodendrocyte Umap cluster (Cluster 0) (Figure 4A), and observed a large proportion of PS19 oligodendrocyte DEGs were reversed in SORLA TG/PS19 hippocampus (Figure 5G), with a high representation of DEGs comprising GO CC “glutamatergic synapse” and “synapse” components (Figure 5H, I). Pathological changes in numerous synapse-related and disease associated (Kenigsbuch, Bost et al. 2022) DEGs in PS19 (PS19 vs WT) oligodendrocytes were reversed in SORLA TG/PS19 hippocampus (Figure 4J). Together, these results indicate that global SORLA upregulation can suppress pathological glial activation and synaptic signatures in PS19 hippocampus.

**Figure 5.**
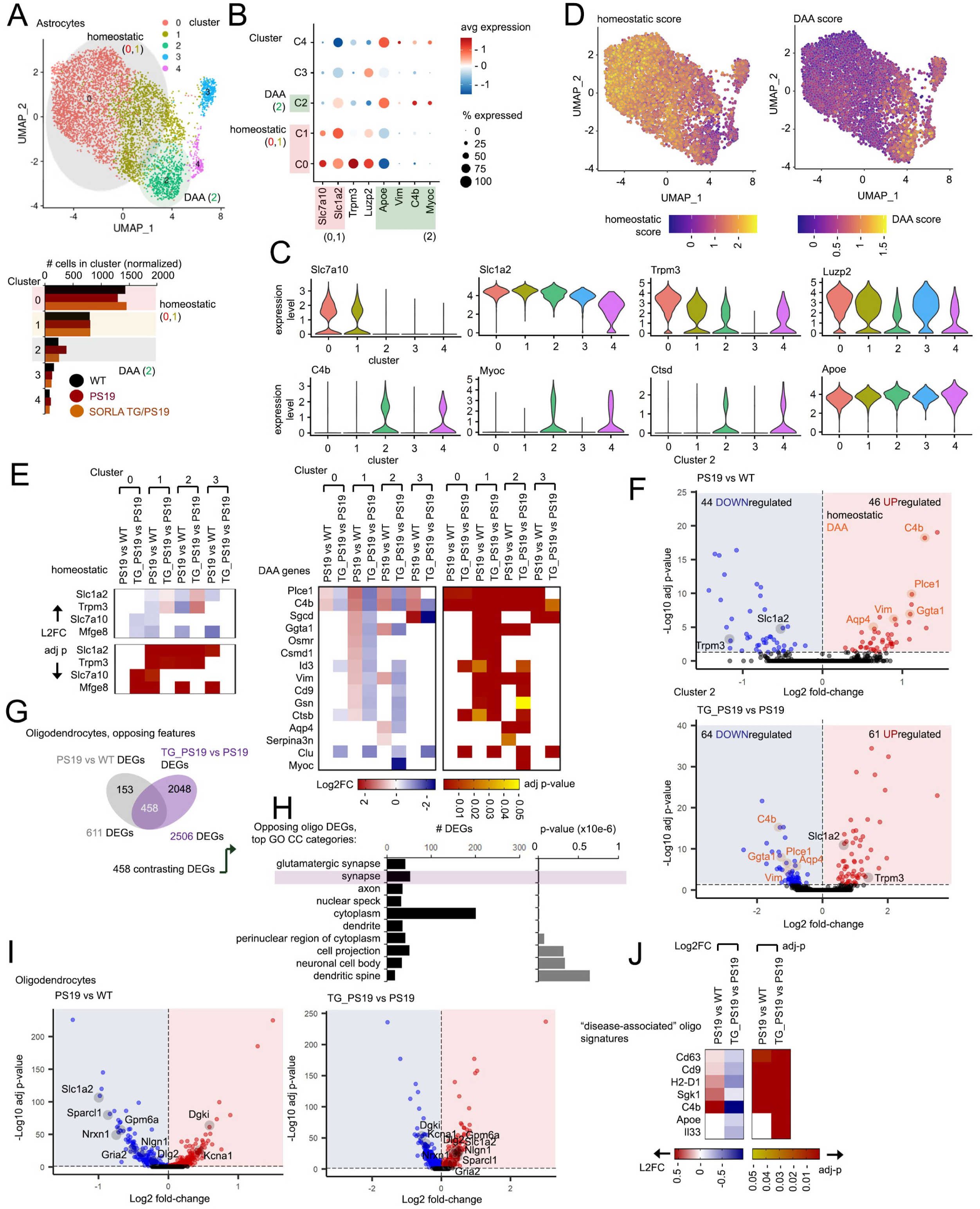
snRNAseq analysis SORLA upregulation suppresses pathological astrocyte and oligodendrocyte signatures in PS19 hippocampus. (A) Umap distribution of 5,579 reclustered astrocyte nuclei. Number of DEGs identified in each cluster in PS19 vs WT and TG_PS19 vs PS19 comparisons are shown in the lower bar graph. (B) Dot plot depicting expression profiles of key astrocyte signatures from each cluster from (A), heat scales for expression levels, as well as % expression (dot size) are shown. (C) Violin plots showing expression profiles of genes indicated in reclustered astrocyte nuclei. (D) Expression of homeostatic and DAA astrocyte genes within reclustered astrocyte transcription profiles. Heat-scaled Umap depicts homeostatic and DAA gene expression scores for reclustered astrocyte nuclei. (E) Expression of homeostatic (left heatmaps) and DAA (right heatmaps) in PS19 vs WT and TG_PS19 vs PS19 comparisons; Log2 fold-change (red/blue) and adj-pvalues (red/yellow) heatmaps depict expression changes and significance for comparisons in the clusters indicated. (F) Volcano plots depicting up and downregulated DEGs in PS19 vs WT and TG_PS19 vs PS19 comparisons in astrocyte Cluster 2 (adj p-value<0.05). Homeostatic (black) and DAA (orange) DEGs featuring opposing expression profiles in PS19 and TG_PS19 astrocytes are indicated. (G, H) Characterizing transcriptomic profiles in oligodendrocytes. (G) Venn diagram depicting opposing DEGs in PS19 oligodendrocytes reversed in SORLA TG hippocampus (TG_PS19 vs PS19). (H) Top 10 GO CC categories observed in 458 opposing DEGs in (G). (I) Volcano plots for DEGs from PS19 vs WT, and TG_PS19 vs PS19 comparisons in oligodendrocytes (adjp<0.05). DEGs related to GO CC “synapse” are indicated. (J) Heatmaps depicting changes in “disease-associated” oligodendrocyte signatures in PS19 vs WT, or TG_PS19 vs PS19 oligodendrocytes; Log2 fold-change (left) and adj-p (right heatmaps) DEG values are shown. See also Figure S4.

### SORLA upregulation can alter transcriptomic and functional response to tau in neurons

Given the importance of neurons in tau pathology, together with a likely role for SORLA in influencing neuronal tau pathology and seeding phenotypes so far, we explored effects of SORLA modulation on neuronal transcriptomic and functional profiles in response to tau. To this end, we characterized snRNA-seq profiles from reclustered neuronal nuclei in WT, PS19 and SORLA TG/PS19 hippocampus. Neuronal snRNA-seq profiles segregated into 11 defined clusters (Figure 6A and Figure S5A), comprising neurons expressing excitatory cell markers such as *Slc17a7* and inhibitory neuronal markers (*Gad1*, *Gad2*) (Figure 6A, B). Clusters 0 and 1 comprised the majority of neuronal DEGs, with markedly fewer DEGs observed in Clusters 2 through 6 (Figure S5B and Table S3); interestingly, a large proportion of PS19 DEGs in Clusters 0/1 were reversed in TG_PS19 neurons (Figure 6C). SORLA upregulation largely reversed GO CC components related to “glutamatergic synapse” DEGs (Figure 6D-F), as well as other components related to neurons, including DEGs related to GO CC “neuron projection”, “dendrite”, and “axons” (Figure 6D). In contrast to upregulation of APOE in PS19 vs WT hippocampus in our proteomics dataset, we observed downregulation of Apoe in PS19 vs WT in neurons, and upregulation of Apoe in SORLA TG/PS19 vs PS19 comparisons by snRNA-seq analysis (cluster 1, Figure S5D, E). We find that suppression of Apoe expression in SORLA TG/PS19 vs PS19 hippocampus is observed in microglia, astrocytes and in oligodendrocytes (Figure S5D, E), suggesting that net Apoe levels may be more heavily influenced by glia rather than neurons in SORLA TG/PS19 and PS19 hippocampus. Together, these results indicate that in addition to glia, SORLA upregulation can also reverse pathological signatures in PS19 neurons.

**Figure 6.**
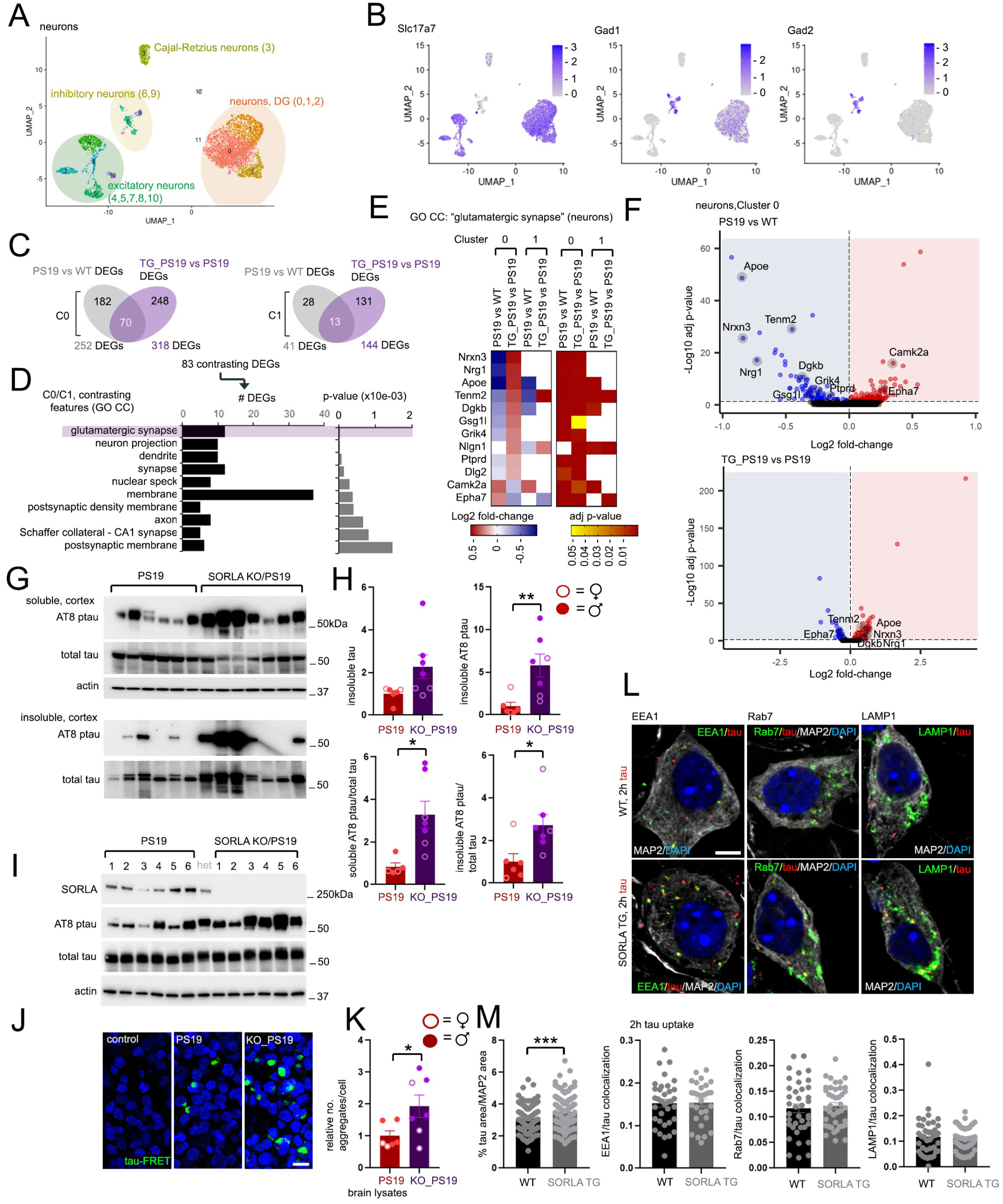
Characterizing neuronal response to tau with SORLA upregulation or deletion. (A) Umap distribution of 5,425 reclustered neuronal nuclei. (B) Expression of Slc17a7 and Gad1, 2 within reclustered neuronal nuclei. Heat-scaled Umap depicts excitatory (Slc17a7) and inhibitory (Gad1, 2) gene expression scores for reclustered neuronal nuclei. (C-E) SORLA upregulation reverses PS19 expression profiles in neuron Clusters 0, 1. (C) Venn diagram showing opposing DEGs (“opposing features”) in PS19 neurons (PS19 vs WT) reversed in SORLA TG hippocampus (TG_PS19 vs PS19) in Cluster 0, 1. (D) GO analysis of 83 contrasting DEGs in PS19 and TG_PS19 neurons from (C), top 10 GO CC categories are shown. (E) Heatmaps depicting Log2 fold-change (left heatmap) and adj p-value (right heatmap) of GO CC “glutamatergic synapse” DEGs identified in (D). (F) Volcano plots depicting up and downregulated DEGs in PS19 vs WT and TG_PS19 vs PS19 comparisons in neuron Cluster 0 (adj p-value<0.05). DEGs featuring opposing expression profiles in PS19 and TG_PS19 neurons are indicated. (G-H) AT8 ptau and total tau from soluble and insoluble fractions in PS19 and SORLA KO/PS19 (KO_PS19) 7 MO cortex. Adjacent graphs depict insoluble tau or AT8 ptau (left graphs) or soluble/insoluble AT8 ptau/total tau ratios (right graphs) normalized to PS19 (set to 1.0). (I-K) (I) Soluble 7 MO PS19 and SORLA KO/PS19 cortical brain extracts were subjected to immunoblot analysis and (J) applied to HEK293 tau-RD cells. (J) Representative tau-FRET images (green) or DAPI (blue) in HEK293 tau-RD cells are shown, bar=20um. (K) Graph depicts relative number of tau-FRET aggregates from PS19 or KO_PS19 animals, normalized to PS19 (set to 1.0) from experiments shown in (J). Only homozygous KO_PS19 samples were plotted for analysis, extracts derived from female or male animals are indicated. (L, M) Characterizing tau uptake in WT and SORLA TG neurons. (L) Representative images of WT or SORLA TG neurons at DIV14 exposed to tau oligomers (15 nM) for 2h, and fixed/stained to visualize EEA1, Rab7 or LAMP1 (green), T13 tau (red) or nuclei (DAPI, blue) colocalization. Neuron area was marked by MAP2 staining (white), bar=5um. (M) Quantification of internalized tau (left graph, % tau area within MAP2 boundaries), or Pearson’s correlation values for tau colocalization with endolysosomal compartments indicated in WT or SORLA TG neurons from images in (L). Plots represent values from one imaged field, and all values were derived from three independent experiments. Graphs depict mean±SE, statistical significance was determined by unpaired Student’s t-test (H, K and M), *p<0.05, **p<0.01, ***p<0.001. See also Figures S5 and S6.

Since SORLA upregulation can attenuate tau pathology and confer neuroprotective effects with respect to synaptic function and transcriptomic features in PS19 brain, SORLA downregulation or dysregulation could potentially exacerbate neuronal tau pathology. To test this, we compared effects of SORLA deletion (SORLA KO) with SORLA upregulation (SORLA TG) on tau pathology and neuron function in PS19 mouse brain and cultured neurons. In contrast to downregulating soluble and insoluble AT8 ptau levels in SORLA TG/PS19 mouse cortex, we found that both soluble and insoluble AT8 ptau/total tau levels were increased in 7MO SORLA KO/PS19 (KO_PS19) cortex compared to PS19 alone (Figure 6G, H). We also found that soluble cortical brain extracts from 7MO SORLA KO/PS19 brain (Figure 6I-K) showed enhanced tau seeding capacity compared to aged-matched PS19 cortex (Figure 6K). Together these results suggest that SORLA deletion can potentially enhance while SORLA upregulation can attenuate pathology in neurons *in vivo*.

SORLA has been previously shown to be an important regulator of endolysosomal trafficking, and downregulation of endosomal retromer components has been observed in AD (Small, Simoes-Spassov et al. 2017). SORLA deletion as well as AD-associated Y1816C, G511R, E270K and Y141C variants can promote endosomal enlargement (Knupp, Mishra et al. 2020, Hung, Tuck et al. 2021, Mishra, Knupp et al. 2023, Jensen, Raska et al. 2024) and dysregulated trafficking of targets such as APP and GLUA1 in iPSC-derived neurons (Mishra, Knupp et al. 2022). Interestingly, pharmacological enhancement of retromer function has been shown to upregulate SORLA and reduce Aβ and tau pathology in a 3xTg AD mouse model (Li, Chiu et al. 2020). Similarly, pharmacological retromer enhancement can reduce Aβ and ptau levels in iPSC-derived neurons with AD-associated SORLA variants (G511R, E270K, Y141C) or deletion/haploinsufficiency (Mishra, Knupp et al. 2023). Given that extracellular tau can be internalized and trafficked to endosomes and lysosomes (Yan and Zheng 2021), we determined whether SORLA modulation can alter tau trafficking and uptake in cultured neurons. We observed significant enhancement of tau internalization in SORLA TG neurons compared to WT following 2h exposure to tau oligomers (Figure 6L, M); we also observed a slight increase in lysosome size in SORLA TG neurons with exposure to tau oligomers (Figure S5F, G).

As a component of the retromer complex, interactions between SORLA and the retromer complex may potentially affect tau pathology/pathogenesis in neurons and other cell types. Although downregulation of retromer components such as VPS35, VPS26 and VPS29 have been observed in combined Aβ/tau (3xTg) secondary mouse models of tauopathy (Li, Chiu et al. 2020), perturbances in retromer in primary tauopathy mouse models remain yet unclear. Our proteomic analysis of PS19 vs WT hippocampus also failed to detect significant dysregulation of the core retromer components (Table S1, Figure S6A). Further, we did not observe alterations in retromer components in our snRNAseq in neurons or glia (Table S3). Proteomics analyses from previous reports indicate that SORLA deletion in iPSC-derived astrocytes, neurons or microglia had little effect on retromer components (Lee, Aylward et al. 2023) (Figure S6B); similarly we find little difference in retromer components (VPS35, VPS26a) in WT compared to SORLA KO, PS19 or SORLA KO/PS19 brain (Figure S6C, D). We observe SORLA in high molecular weight complexes (∼660kDa) by SEC fractionation in mouse brain which appear to be unchanged in SORLA TG, PS19 and SORLA TG/PS19 mice (Figure S6E, F). We also observe VPS35 and VPS26a in a complex of ∼450kDa in WT mouse brain with little to no variation in SORLA TG, KO, PS19 or TG_PS19, KO_PS19 backgrounds (Figure S6F). We also observe no significant variation in SORLA or retromer levels (VPS35, VPS26a, VPS26b) in cultured WT vs PS19 neurons (Figure S6G, H).

SORLA upregulation could attenuate neurogenerative phenotypes such as brain atrophy, as well as various transcripts related to synaptic proteins, indicating that SORLA upregulation has strong neuroprotective effects in neurons. Indeed, SORLA has been previously regulate and traffic neuronal components, including sorting/trafficking the GDNF/GFRa1 complex (Glerup, Lume et al. 2013), regulating calpain-dependent degradation of synapsin (Hartl, Nebrich et al. 2016), and trafficking glutamate receptors such as GLUA1 (Mishra, Knupp et al. 2022), which could consequently affect neurodegenerative outcome. We also further characterized synaptic targets potentially perturbed in neurons and glia in PS19 hippocampus. We observed impaired expression of *Nlgn1*, a postsynaptic cell surface factor essential for maturation and organization of synaptic junctions (Sudhof 2008, Sindi, Tannenberg et al. 2014), in both PS19 neurons (Figure S5C) and microglia (Figure S6I) was restored in SORLA TG/PS19 hippocampus. *Nlgn1-Nrxn1* interaction stabilize and facilitate proper function of synaptic junctions (Sudhof 2008). Interestingly, *Nrxn1* was downregulated in all PS19 glial cell types and upregulated in SORLA TG/PS19 astrocytes and oligodendrocytes (Figure S6I). Although a role for *Nrxn1* as a mediator of synaptic signaling is well established in neurons (Sudhof 2008), *Nrxn1* expression, function and dysregulation in glial cell types with tau proteotoxicity has yet to be described. We characterized *Nrxn1* transcript levels by RNAscope analysis in astrocytes, neurons and microglia (Figure S6J). Using antibodies to identify astrocytes (GFAP), neurons (NeuN) or microglia (Iba1), we quantified the number of *Nrxn1* puncta specifically in these cell types in 9MO WT, SORLA TG, PS19 or SORLA TG/PS19 (PS19/TG) hippocampus (CA3) (Figure S6J, K). We observed significantly reduced *Nrxn1* puncta in astrocytes and microglia, as well as a trending decrease in Nrxn1 puncta in neurons in PS19 hippocampus; we observed significant elevation of *Nrxn1* puncta in neurons and trending elevations in *Nrxn1* puncta in PS19/TG compared to PS19 hippocampus (Figure S6K).

Together, our results indicate that SORLA upregulation can reverse neuronal signatures and targets in PS19 hippocampus, and enhance tau uptake and trafficking to lysosomes in neurons. However, retromer levels are not significantly perturbed in PS19 neurons or PS19 mouse brain.

### Pathological alterations in cell-cell communication/signaling networks are restored with SORLA upregulation

Given that SORLA upregulation can reverse transcriptomic changes in various cell types in PS19 hippocampus, we explored whether specific cell-cell interaction networks could be potentially restored in SORLA TG/PS19 animals. Using the CellPhoneDB database (v5.0.0) to infer cell-cell communication networks from our snRNA-seq analyses (Efremova, Vento-Tormo et al. 2020), we identified ligand/receptor complexes enriched in PS19 (pathogenic) or WT/TG_PS19 (non-pathogenic) genotypes (Figure 7A and Figure S6L). As expected, we also observed enrichment of SORLA (*Sorl1*)/APP cell-cell interaction pairs in WT and TG_PS19 astrocyte/neurons (Figure 7A).

**Figure 7.**
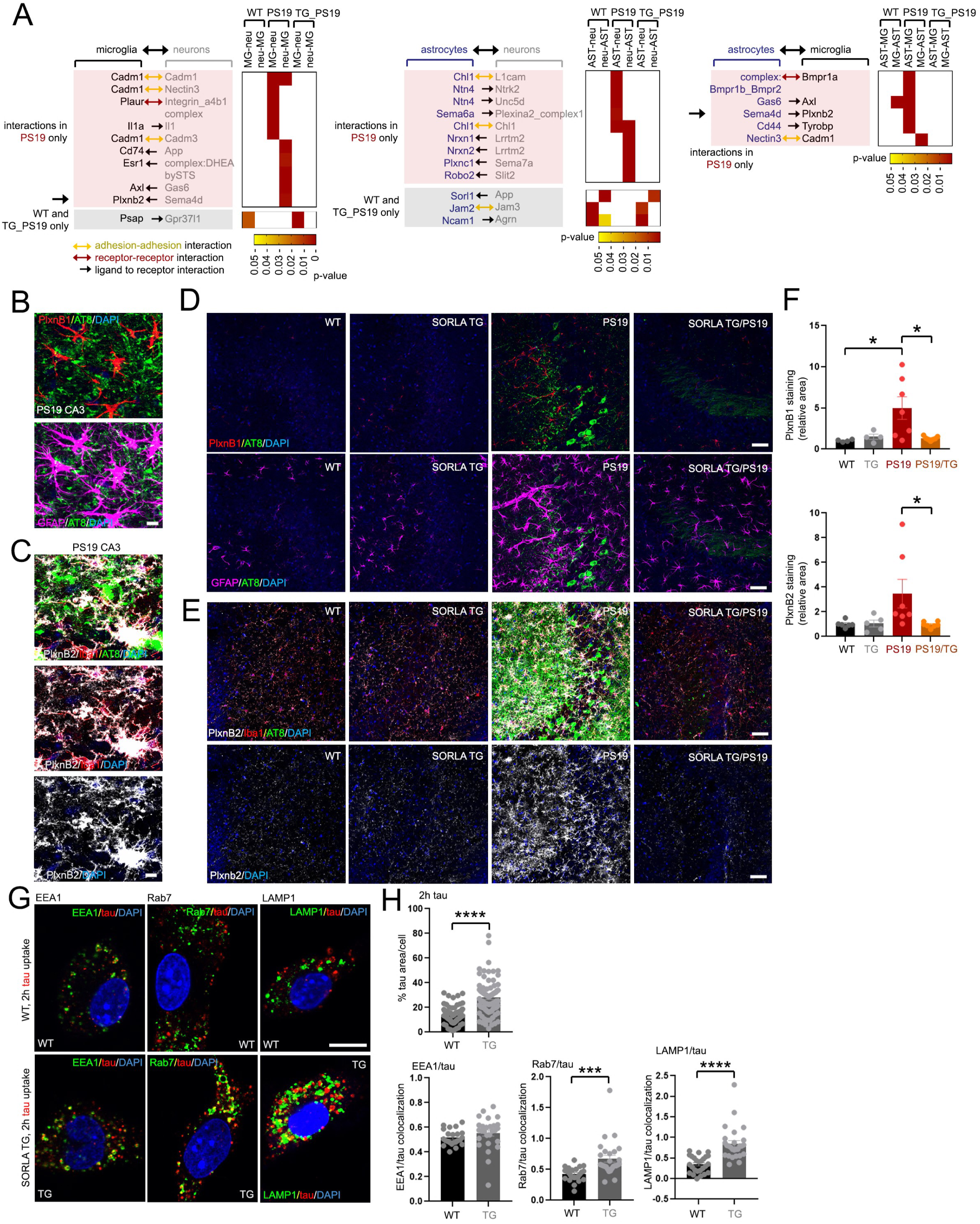
Characterizing changes in receptor/ligand pairs in PS19 hippocampus with SORLA upregulation. (A) Identification of changes in ligand/receptor pairs in microglia/neurons (left), astrocytes/neurons (middle) and astrocytes/microglia (right) using CellPhoneDB analysis of snRNAseq profiles in WT, PS19 and SORLA TG/PS19 (TG_PS19) hippocampus. Changes in matched receptor/ligand pairs specific to PS19 are highlighted in red, paired interactions specific to WT and TG_PS19 hippocampus are highlighted in gray. Adjacent heatmaps depict p-values calculated for each receptor/ligand interaction (replicate p-values are averaged) in paired cell types. (B-F) Quantification of PlxnB1 and PlxnB2 in 9MO WT, SORLA TG, PS19 and SORLA TG/PS19 hippocampus (CA3). (B) Co-localization of PlxnB1 (red) and AT8 p-tau (green) (upper panel) and GFAP (lavender) and AT8 (green) (lower panel) in PS19 hippocampus (CA3). Bar = 10um. (C) Co-localization of PlxnB2 (white), Iba1 (red), and AT8 p-tau (green) (upper panel) in PS19 hippocampus (CA3). Merged PlxnB2/Iba1 (middle) and PlxnB2 alone (lower panel) is shown. Bar = 10um. (D) Representative images of PlxnB1 (red), AT8 p-tau (green) (upper panels) or GFAP (lavender) and AT8 p-tau (green) (lower panels) in 9MO CA3 hippocampus. Bar = 50um. (E) Representative images of PlxnB2 (white), Iba1 (red), and AT8 p-tau (green) (upper panels) or Plxnb2 (white) (lower panels) in 9MO CA3 hippocampus. Bar = 50um. (F) Quantification of Plxnb1 (upper graph) (WT: 2F, 2M; STG: 2F, 2M; PS19: 4F, 3M; STG/PS19: 4F, 3M.) and Plxnb2 (lower graph) (WT: 3F, 3M; STG: 3F, 2M; PS19: 4F, 3M; STG/PS19: 4F, 3M) staining (relative area) from representative images in (D) and (E) relative to WT (set to 1.0). Individual plots represent averaged quantified area measurements in individual animals. Graphs represent mean±SE; statistical significance was determined by One-way ANOVA (F), *p<0.05, ***p<0.001, ****p<0.0001. See also Figure S6.

We observed enrichment of semaphorin signaling ligands (*Sema6a* – astrocytes, *Sema4d* – neurons, astrocytes) and Semaphorin receptors (*Plexina2* – neurons, *Plxnb2* – microglia) specifically in PS19 hippocampus (Figure 7A). The Semaphorin protein family comprises an extracellular Sema domain as well as a Plexin-Semaphorin-Integrin domain required for mediating homotypic and heterotypic interactions (Zhou, Gunput et al. 2008), (Carulli, de Winter et al. 2021). Interestingly, SEMA4D levels are upregulated in human Huntington’s disease (HD) and AD brain (Evans, Mishra et al. 2022); while PlxnB1 expression correlates with Aβ pathology and cognitive impairment in AD (Mostafavi, Gaiteri et al. 2018). Since antibody-based Sema4D targeting is shown to reverse cognitive impairment and astrogliosis in AD mice (Evans, Mishra et al. 2022), we explored whether Sema4D and cognate PlxnB1 and PlxnB2 receptors were altered in PS19 hippocampus.

*Sema4d* expression is enriched in mouse oligodendrocytes and microglia (Moreau-Fauvarque, Kumanogoh et al. 2003, Bennett, Bennett et al. 2016). We observed a variable, but slight increase in Sema4D levels in PS19 hippocampus by immunoblot, and comparatively reduced Sema4D levels in SORLA TG/PS19 animals (Figure S7A, B). PlxnB1 binds Sema4D with high affinity (Tamagnone, Artigiani et al. 1999) and is highly expressed in mouse astrocytes (Bennett, Bennett et al. 2016), whereas PlxnB2 is expressed primarily in microglia (Bennett, Bennett et al. 2016). We observed high expression of PlxnB1 in astrocytes (Figure 7B) and Plxnb2 in microglia (Figure 7C). Basal PlxnB1 and PlxnB2 levels were quite low in WT and SORLA TG hippocampus (CA3), and markedly increased in PS19 animals (Figure 7D-F). Remarkably, SORLA upregulation significantly suppressed PlxnB1 and PlxnB2 levels in PS19 hippocampus (Figure 7D-F). Together, these results demonstrate that SORLA upregulation can alter pathological PlxnB1/PlxnB2 induction in PS19 hippocampus.

### SORLA deletion exacerbates pathogenic changes in PS19 brain

Reduced SORLA levels have been reported in AD patient lymphoblasts (Scherzer, Offe et al. 2004) and brain (Andersen, Reiche et al. 2005), indicating that SORLA loss of function may be linked to AD pathogenesis. Because our results suggest that SORLA upregulation can suppress expression of mediators of tau proteotoxicity, we further explored whether SORLA deletion could alter pathogenic effects in PS19 mouse brain. We generated homozygous SORLA KO mice in a PS19 background (“KO_PS19”) and compared Sema4D levels in WT, homozygous SORLA KO (“KO”), PS19, and heterozygous (“het_PS19”) and KO_PS19 deletion strains (Figure S7C, D). We observed increased Sema4D levels in KO_PS19 vs PS19 by immunoblot, with no significant change in het_PS19 over PS19 in 9MO hippocampus (Figure S7C, D). To further characterize changes in astrocytes and microglia in KO_PS19 hippocampus compared to PS19 or KO alone, we sampled regions of interest (ROIs) in CA1, CA3 and DG subregions within the hippocampus in KO_PS19, PS19 and KO animals (Figure 8A), and characterized transcriptomic changes associated with astrocytes (GFAP staining) and microglia (Iba1 staining). We observe greater numbers of astrocyte DEGs in KO_PS19 vs PS19 (945 DEGs) compared to KO_PS19 vs KO (145 DEGs) (Figure S7E, Table S4) indicating that SORLA deletion can enact dramatic changes in PS19 astrocytes. We find that SORLA deletion can increase expression of astrocyte DAA DEGs over PS19 (KO_PS19 vs PS19; *Aqp4, Clu, Serpina3n*, *Ctsb*) as well as KO (KO_PS19 vs KO; *Aqp4, Clu, Serpina3n*, *C4b*) (Figure 8B, C and Figure S7F; Table S4), indicating that SORLA deletion can aggravate astrocyte pathology in aged PS19 brain. GO analysis of KO_PS19 vs PS19 DEGs yield Biological Process (BP) categories related to transcription, cell differentiation, and cytokine production/innate immune response (Figure S7G).

**Figure 8.**
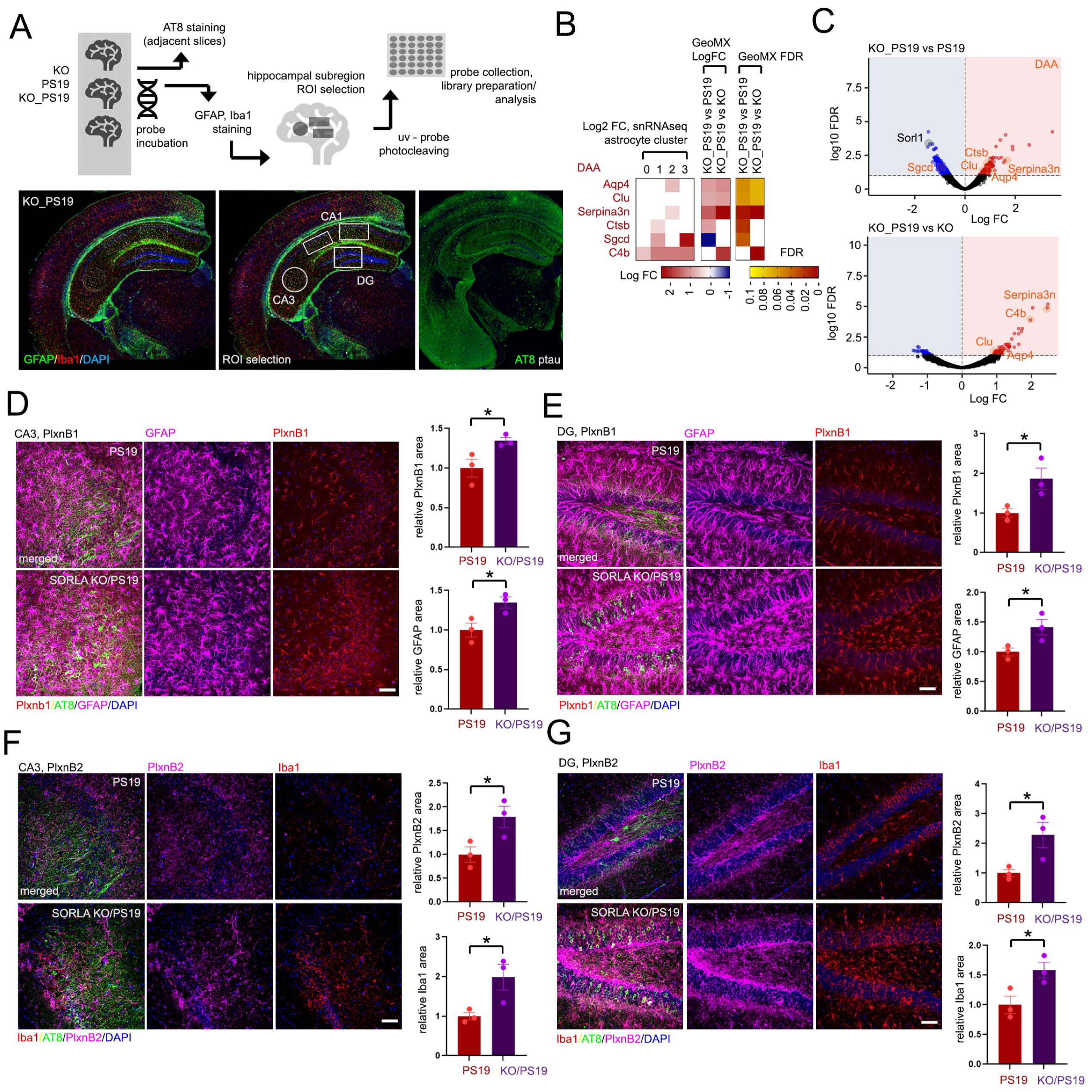
SORLA deletion exacerbates astrocyte and microglia activation in PS19 hippocampus. (A) Schematic of the GeoMx workflow; processing and analysis of SORLA KO, PS19 and SORLA KO/PS19 (“KO_PS19”) coronal sections (upper panel). Lower panels: representative hemisection of 9MO KO_PS19 hippocampus stained for GFAP (green) Iba1 (red) and nuclei (DAPI, blue), processed for GeoMX analysis. ROI subregions sampled are indicated (lower, middle panel), and AT8 ptau staining is shown in an adjacent hemisection (green, lower, right panel). (B) Changes in DAA astrocyte DEGs identified in PS19 vs WT animals by snRNAseq, and in KO_PS19 vs PS19 or KO_PS19 vs KO comparisons by GeoMX analysis. Heat scales for Log fold-change or FDR are shown. (C) Volcano plots showing DEGs identified in KO_PS19 vs PS19 (top) or KO_PS19 vs KO (bottom) comparisons by GeoMX analysis. DAA DEGs are indicated in orange (FDR<0.1). (D-F) SORLA deletion upregulates Plxnb1 and Plxnb2 levels in PS19 hippocampus. Representative images and quantification of PlxnB1 (D, E) and PlxnB2 (F, G) in CA3 (D, F) and DG (E, G) from 9MO PS19 and SORLA KO/PS19 hippocampus as indicated. Hippocampus stained for PlxnB1 (red) is also counterstained for AT8 ptau (green), GFAP (purple) and nuclei (DAPI, blue); hippocampus stained for PlxnB2 (purple) is also counterstained for Iba1 (red), AT8 ptau (green) and nuclei (DAPI, blue) (bar = 50um). Adjacent graphs depict relative PlxnB1, GFAP, PlxnB2 and Iba1 staining area from PS19 and SORLA KO/PS19 (“KO/PS19”) in hippocampal CA3 (D, F) or DG (E, G) regions quantified from 9MO animals; plots represent quantified measurements from individual animals, normalized to PS19 (set to 1.0). (PS19: 2F, 1M; SKO/PS19: 3M). Graphs represent mean±SE; statistical significance was determined by unpaired Student’s t-test (D, E), *p<0.05. See also Figures S7 and S8.

In microglia, we observed more KO_PS19 vs KO DEGs compared to KO_PS19 vs PS19 (Figure S7I, Table S4), and DAM signatures showed significant upregulation in KO_PS19 hippocampus compared to KO (Figure S7J-L). We also performed GeoMx analysis on NeuN-stained neurons in KO, PS19 and KO_PS19 hippocampus (Figure S8A), and similar to microglia we observe the highest change in KO_PS19 vs KO comparisons with relatively few DEGS in KO_PS19 vs PS19 comparisons (Figure S8B, C and Table S4). Remarkably, we find that KO_PS19 vs KO DEGs were almost exclusively upregulated (1582 upregulated, 2 downregulated) (Figure S8C), indicating that PS19 has strong effects in upregulating transcripts in KO neurons. We also observed general upregulation of both Sema4D and PlxnB2 in microglia in KO_PS19 hippocampus compared to PS19 and control (Figure S8D). Interestingly, PlxnB1/B2 and Sema4D levels have been shown to be upregulated in aged thy-TAU22 mice by bulk RNAseq analysis (Sierksma, Lu et al. 2020) and in PlxnB1 in human GRN-FTD astrocytes (Gerrits, Giannini et al. 2022) (Figure S8E). Since we observed marked upregulation of PlxnB1 and PlxnB2 in PS19 hippocampus compared to WT in the hippocampal CA3 region which was suppressed in SORLA TG/PS19 animals (Figure 7B, C), we determined whether SORLA upregulation can exacerbate PlxnB1 and PlxnB2 expression. We found PlxnB1 levels in SORLA KO hippocampus comparable to WT, and appeared to be elevated in PS19 DG region (Figure S8F, G). We observed a significant increase in PlxnB1 and PlxnB2 levels in CA3 and DG (Figure 8D-G, and Figure S8F, G) regions in SORLA KO/PS19 as well as GFAP and Iba1 staining respectively compared to PS19 alone. This suggests that while SORLA upregulation can suppress PlxnB1/B2 levels, SORLA deletion can exacerbate PlxnB1/B2 expression.

## Discussion

Although numerous genetic risk factors have been identified in sporadic AD, commonalities between various risk factors such as APOE and various SORLA variants remain somewhat unclear. In the case of APOE, dysregulation of these AD risk factors affect both Aβ and tau pathogenesis in various mouse models. Murine *Apoe* deletion reduced Aβ deposition in various transgenic APP mouse models (Bales, Verina et al. 1997, Bales, Verina et al. 1999, Holtzman, Fagan et al. 2000, Irizarry, Cheung et al. 2000, Ulrich, Ulland et al. 2018), and expression of human APOE4 is seen to exacerbate pathological features including tau pathology, ventricle dilation and glial activation (Shi, Yamada et al. 2017). Removal of the neuronal APOE4 transgene in APOE4/PS19 mice attenuates tau pathology, ventricle dilation, gliosis and neuronal hyperexcitability (Koutsodendris, Blumenfeld et al. 2023), indicating that APOE is a mediator of both Aβ and tau pathogenesis. Retromer deficiency or dysregulation which has been observed in AD brain (Small, Kent et al. 2005) can also perturb APOE levels, for example neuronal Vps35 deletion can elevate APOE in mouse CSF (Simoes, Neufeld et al. 2020) as well as hippocampus, likely through glial activation observed with neuronal Vps35 deletion (Qureshi, Berman et al. 2022). Although a role for SORLA has been well established in trafficking APP (Andersen, Reiche et al. 2005, Fjorback, Seaman et al. 2012, Huang, Zhao et al. 2016), Aβ uptake and clearance in cultured cells (Caglayan, Takagi-Niidome et al. 2014) as well as ESC-derived microglia models (Liu, Zhu et al. 2020), how SORLA can affect tau remains relatively unexplored. Interestingly, SORLA haploinsufficiency in minipigs results in elevated Aβ and tau CSF levels with little change in brain Aβ/tau pathology (Andersen, Bogh et al. 2022). This suggests that in addition to trafficking and internalizing Aβ (Caglayan, Takagi-Niidome et al. 2014, Liu, Zhu et al. 2020), SORLA may also potentially affect tau accumulation.

In agreement with this notion, recent evidence suggests that SORLA can interact directly with tau through interactions with the N-terminal SORLA VSP10 domain, and SORLA can promote tau aggregation and seeding from tau-enriched fractions in AD brain samples in an HEK293 tau-RD FRET biosensor model (Cooper, Lathuiliere et al. 2024), suggesting that SORLA can affect cytosolic seeding of tau internalized from an extracellular source. While observations that SORLA can potentially enhance intracellular tau seeding from exogenous tau exposure are interesting, these effects are so far only seen in cultured cell lines (Cooper, Lathuiliere et al. 2024). Our results here indicate that SORLA upregulation can attenuate, while SORLA deletion can aggravate pathological tau phosphorylation and seeding in aged PS19 mouse brain, where tau pathology resides mostly within neurons. This suggests that short-term modulation of SORLA in a human cell line may have differing effects on exogenously internalized tau (Cooper, Lathuiliere et al. 2024) compared to global SORLA modulation in aged PS19 mouse brain. Given that tau pathology originates in neurons, effects of SORLA in suppressing tau phosphorylation and aggregation in neurons may be key to attenuating pathogenic effects related to tau. This may trigger other events related to tau pathogenesis and toxicity; for example, SORLA upregulation can suppress induction of key drivers of tau pathogenesis such as C1q (Dejanovic, Huntley et al. 2018, Dejanovic, Wu et al. 2022) and Apoe (Shi, Manis et al. 2019). In support of the notion that SORLA may be in fact protective with respect to tau pathology/pathogenesis, recent work indicates that SORLA deletion in iPSC-derived neurons enhances tau phosphorylation (Lee, Aylward et al. 2023). Since SORLA expression is thought to be reduced (Scherzer, Offe et al. 2004, Andersen, Reiche et al. 2005) or downregulated by AD-associated stop-gain variants (Pottier, Hannequin et al. 2012, Nicolas, Charbonnier et al. 2016, Holstege, van der Lee et al. 2017), it is likely that SORLA dysfunction is deleterious to tau pathogenesis.

Activation of microglia and astrocytes is a common feature of AD and other tauopathies (Di Benedetto, Burgaletto et al. 2022, Chen and Yu 2023, Deng, Wu et al. 2024); characterization of transcriptional signatures in mouse models of AD and tauopathy have profiled distinctive pathological states in microglia and astrocytes states. Interestingly, many of the activated “disease-associated microglia” (DAM) signatures observed in 5xFAD microglia (Keren-Shaul, Spinrad et al. 2017) are also observed in other models of neurodegeneration (neurodegenerative microglia, MGnD) including amyotrophic lateral sclerosis (ALS), or multiple sclerosis (MS) (Krasemann, Madore et al. 2017). Signatures associated with microglia DAM states as well as MHC-related transcripts are also observed in PS19 mouse models (Udeochu, Amin et al. 2023, Wang, Martini-Stoica et al. 2024). Astrocytes also feature distinctive DAA signatures in 5xFAD mice (Habib, McCabe et al. 2020) that are also altered in PS19 brain (Koutsodendris, Blumenfeld et al. 2023). Together, these results indicate that microglia and astrocytes feature overlapping transcriptomic signatures that coincide with activation in Aβ or tau mouse models of neurodegeneration and proteotoxicity.

Tau has been shown to be associated with synaptic dysfunction, and while tau is normally associated with and stabilizes microtubules, tau phosphorylation results in dissociation from microtubules, tau redistribution to somatodendritic compartments and synaptic dysfunction (Hoover, Reed et al. 2010, Noble, Hanger et al. 2013, Spires-Jones and Hyman 2014). Interestingly, reducing tau levels can attenuate synaptic impairment in hAPP (Roberson, Scearce-Levie et al. 2007) and APP23 (Ittner, Ke et al. 2010) mouse models, indicating that tau is important in driving toxic effects associated with Aβ proteotoxicity. Given that tau pathology predominantly resides within neurons, we consider that SORLA upregulation has a particularly strong effect in limiting tau pathology in neurons. Previous groups have reported that SORLA deletion results in endosomal (Knupp, Mishra et al. 2020, Hung, Tuck et al. 2021), as well as lysosomal defects (Hung, Tuck et al. 2021) in iPSC-derived neurons. Our results indicate that SORLA upregulation enhances tau uptake in both neurons and microglia, and increases tau trafficking to late endosomes/lysosomes in microglia. While it is not possible to specifically determine how cell-specific modulation of SORLA can affect tau using the global transgenic overexpression/deletion models used here, we are interested in further characterizing specific effects of SORLA on tau in neurons, and the extent of SORLA modulation on glia in influencing overall tau pathology.

Our results also indicate that SORLA upregulation could affect synaptic integrity and function through mechanisms associated with synaptic stability and/or elimination. In addition to being key drivers of tau pathogenesis in PS19 brain, APOE and C1q have been shown to be important mediators of synaptic function. *Apoe* deletion in mice results in synaptic loss and cognitive impairment, which could be restored by peripheral Apoe expression (Lane-Donovan, Wong et al. 2016). Previous studies have shown that expression of human APOE4 can enhance C1q accumulation and increase astrocyte-mediated synaptic pruning in mouse brain (Chung, Verghese et al. 2016); similarly, iPSC-derived APOE4/E4 organoids feature enhanced apoptosis and impaired synaptic integrity compared to APOE3/E3 organoids (Zhao, Fu et al. 2020). In addition to effects observed with APOE4 in enhancing C1q accumulation in mouse brain (Chung, Verghese et al. 2016), APOE can bind to activated C1q and inhibit activation of the classical complement cascade (Yin, Ackermann et al. 2019). APOE/C1q complexes also correlated with cognitive impairment and atherosclerosis in the choroid plexus in mouse and human AD brain, implicating that disease pathogenesis may involve in direct interactions between APOE and C1q (Yin, Ackermann et al. 2019). C1q is a component of the classical complement pathway, and has been shown to be important for tagging synapses for microglial-mediated uptake and elimination in the retinogeniculate early during development (Stevens, Allen et al. 2007). In AD, C1q is also involved in pathologically labeling synapses for microglial elimination in response to Aβ proteotoxicity in J20 (hAPP) (Hong, Beja-Glasser et al. 2016), as well as PS19 mice (Zhong, Sheng et al. 2023), as well as astrocyte-mediated synapse elimination in PS19 mouse brain (Dejanovic, Wu et al. 2022). C1q expression in microglia is also an important component of astrocyte activation, with implications for microglia/astrocyte crosstalk in AD (Liddelow, Guttenplan et al. 2017). Our results that SORLA upregulation can attenuate expression of both APOE and C1q indicates that SORLA can attenuate a spectrum of pathogenic effects associated with these components ranging from synaptic uptake to neuroinflammatory glial activation.

In addition to APOE and C1q, we identified modulation of other potential key targets or pathogenic enhancers of tau toxicity with SORLA upregulation. Numerous synaptic components dysregulated in PS19 hippocampus was found to be restored by proteomic and snRNA-seq analysis, suggesting that SORLA upregulation in neurons likely has strong cell autonomous effects in restoring components related to synaptic function in neurons. For example, we observed downregulation of the synaptic cell-adhesion receptor, Nrxn1 in PS19 hippocampus across glia and neuronal cell types which was restored with SORLA upregulation. Neurexins at neuronal presynaptic terminals typically interact with neuroligins at post-synaptic densities, and mediate downstream intracellular signaling cascades through a C-terminal class II PDZ motif through interactions with the MAGUK protein kinase CASK (Hata, Butz et al. 1996) and protein 4.1 (Biederer and Sudhof 2001). In addition to mediating synaptic function at neuronal junctions, neurexin/neuroligin interactions have been observed in neurons/astrocyte interactions potentially mediating astrocyte morphology and synaptogenesis (Stogsdill, Ramirez et al. 2017). Interestingly, neurexins can interact with Aβ oligomers to potentially interfere with presynaptic differentiation in reconstituted synapse assays (Naito, Tanabe et al. 2017), and blocking Aβ/neurexin interactions using anti-neurexin antibodies can potentially reverse Aβ-mediated synapse loss in cultured neurons and memory impairment in mice (Brito-Moreira, Lourenco et al. 2017). Expression of β- neurexins are also observed to decrease in J20 AD mouse brain (Naito, Tanabe et al. 2017); Nrxn1 was also seen to be downregulated in human AD astrocytes (Lau, Cao et al. 2020). Our observations here indicate that neurexins, namely Nrxn1 is also downregulated in response to tau proteotoxicity in PS19 hippocampus over glial and neuronal cell types, and its expression is restored with SORLA upregulation.

We also observed changes in pathological ligand/receptor pairs in PS19 hippocampus that were attenuated with SORLA upregulation. We were particularly interested in potential changes in the semaphorin family of extracellular ligands and plexin-family of semaphorin receptors. PlxnB1 correlates with Aβ pathology and cognitive impairment in human AD brain (Mostafavi, Gaiteri et al. 2018) and Sema4d is a potential driver of astrogliosis in AD mouse brain (Evans, Mishra et al. 2022). Plxnb1 levels are also associated with increased Aβ load and clinical degenerative features by targeted proteomic analysis in human AD brain (Yu, Petyuk et al. 2018), and can regulate microglia/astrocyte distancing and activation, where Plxnb1 deletion can enhance plaque compaction and reduce neuroinflammation in AD (APP/PS1) mouse brain (Huang, Wang et al. 2024). Sema4d/PlxnB2 interactions were also implicated in glial activation in response to EAE demyelination models of MS (Clark, Gutierrez-Vazquez et al. 2021). Our observations that Plxnb1 and Plxnb2 are both highly upregulated in astrocytes and microglia in PS19 respectively indicate that semaphorin/plexin signaling is potentially a tau-dependent driver of glial activation normalized with SORLA upregulation. As our results strongly indicate that SORLA upregulation can attenuate both microglia and astrocyte activation, it may be interesting to determine in future studies whether attenuation of APOE/C1q and semaphorin/plexin individually or coordinately contribute to pathogenic phenotypes in PS19 mouse brain, and explore relationships between SORLA expression and pathogenic APOE/C1q and semaphorin/plexin in human AD and tauopathy brain.

Here, we have established pleiotropic effects in reversing features associated with tau toxicity in PS19 mouse brain with global SORLA upregulation. Although we are unable to clearly distinguish functional cell-specific effects associated with transgenic SORLA expression using this model, SORLA is expressed in a variety of CNS cell types, and pathological effects due to AD stop-gain variants are likely derived from combined effects from glia, neurons and functional cross-talk between various CNS cell types. Although the P301S tauopathy model used here features varied, yet strong pathological effects and is widely characterized in the AD/tauopathy field, humanized tau models such as the hTau model comprising a P1-derived artificial chromosome (PAC) copy of the human *MAPT* gene locus in a murine *Mapt* KO background (Duff, Knight et al. 2000, Tucker, Meyer et al. 2001, Andorfer, Kress et al. 2003) also feature age-dependent tau pathology, neuronal loss and memory deficits (Andorfer, Kress et al. 2003, Andorfer, Acker et al. 2005, Polydoro, Acker et al. 2009). The hTau-KI human *MAPT* knock-in mouse model lacks apparent tau pathology, but has been shown to enhance tau propagation with exposure to AD-derived tau aggregates (Hashimoto, Matsuba et al. 2019, Saito, Mihira et al. 2019). Given that SORLA is an AD variant, and AD onset largely occurs independently of *MAPT* tau variants, whether our observations here can be also reflected in other humanized tauopathy mouse models would be interesting. We also note that use of the PS19 tauopathy model only features tau pathology in the absence of Aβ pathology; transgenic or humanized APP models may be needed to better recapitulate neurodegenerative effects associated with SORLA modulation in AD. We anticipate future studies using humanized MAPT, and/or Aβ/tau models in combination, as well as cell-specific SORLA KO models will allow us to further distinguish and characterize cell-specific effects associated with SORLA dysregulation in AD and related tauopathies.

## ACKNOWLEDGMENTS

We dedicate this article to the memory of Dr. Huaxi Xu. This work was supported by grants from National Institute of Health (R01 AG061875, RF1 AG070391 and R01 AG085498 to T.Y.H.). We thank all the members in Huang lab for the critical discussion. Special thanks to Svetlana Maurya and Valesca Anschau (SBP Proteomics Core), and Guillermina Garcia (SBP Histopathology Core) for technical assistance. We are grateful to have the support from the Sanford Burnham Prebys Animal Resources Core, Histology Core, Cancer Metabolism Core and Genomics Core support. SBP core services were supported through a Sanford Burnham Prebys NCI Cancer Center Support Grant (P30 CA030199). This publication includes data generated at the UC San Diego IGM Genomics Center utilizing an Illumina NovaSeq 6000 that was purchased with funding from a National Institutes of Health SIG grant (#S10 OD026929), and GeoMx was performed on a DSP instrument obtained from NIH funding at SBP (S10 OD030285).

## CONTRIBUTIONS

H.H. and T.Y.H. designed the study and wrote the manuscript. H.H. performed most experiments and analyzed the data, assisted by W.Y., J.C. and P.S.; J.B. and J.P-C. performed electrophysiology analysis. R.A.P. performed GeoMx library preparation and sequencing. A.C. performed bioinformatic analysis for the proteomics dataset. R.M. processed the snRNA-seq raw data, C.H.S. and R.M. performed quality control/doublet removal procedures, and GeoMx raw data and generated the related plots for the figures. T.Y.H., J.C., R.M. and C.H.S. performed downstream bioinformatic for the “-omic” datasets, C.H.S. performed CellPhoneDB analysis, and K.Y.Y. provided input supervising data analysis and interpretation. A.H., W.Y., S.F. performed Size Exclusion Chromatography experiments. Q.X. conducted blind analysis of cell culture images using AI-driven image processing with Zeiss Arivis software. T.Z. provided critical discussion, insight and advice. T.Y.H. supervised the research, and provided guidance throughout the study.

### Declaration of interests

All the authors declare no competing interests.

## CONTRIBUTIONS

A. H. and T.Y.H. designed the study and wrote the manuscript. H.H. performed most experiments and analyzed the data, assisted by W.Y., J.C. and P.S.; J.B. and J.P-C. performed electrophysiology analysis. R.A.P. performed GeoMx library preparation and sequencing. A.C. performed bioinformatic analysis for the proteomics dataset. R.M. processed the snRNA-seq raw data, C.H.S. and R.M. performed quality control/doublet removal procedures, and GeoMx raw data and generated the related plots for the figures. T.Y.H., J.C., R.M. and C.H.S. performed downstream bioinformatic for the “-omic” datasets, C.H.S. performed CellPhoneDB analysis, and K.Y.Y. provided input supervising data analysis and interpretation. A.H., W.Y., S.F. performed Size Exclusion Chromatography experiments. Q.X. conducted blind analysis of cell culture images using AI-driven image processing with Zeiss Arivis software. T.Z. provided critical discussion, insight and advice. T.Y.H. supervised the research, and provided guidance throughout the study.

## METHODS

### RESOURCE AVAILABILITY

#### Lead contact

Further information and requests for resources and reagents should be directed to and will be fulfilled by the Lead Contact, Timothy. Y. Huang.

#### Materials availability

This study did not generate new unique reagents.

#### Data and code availability

All sequencing data (snRNA-seq, GeoMx) described in this study has been deposited in the NCBI Gene Expression Omnibus (GEO) under the following SuperSeries accession number GSE278216: the GeoMx dataset has been assigned the accession no. GSE278067, and the snRNA-seq dataset has been assigned the accession no. GSE278215. The data can be accessed through the following anonymous link: https://www.ncbi.nlm.nih.gov/geo/query/acc.cgi?acc=GSE278216, using the entry token “wlmrgygmdpwjzkl”. Proteomics raw and search data included in this study are deposited in the ProteomeXchange database under the project accession no. PXD056276. Other data or codes will be made available upon request.

### EXPERIMENTAL MODEL AND SUBJECT DETAILS

#### Mouse lines

SORLA-Rosa26 (SORLA TG) mouse lines overexpressing SORLA through a CMV/β-actin promoter element was previously established (Caglayan et al., 2014). SORLA KO mice were obtained from Thomas Willnow (Max Delbruck Center for Molecular Medicine, Berlin) (Andersen, Reiche et al. 2005). PS19 mouse lines (B6;C3- Tg(Prnp-MAPT*P301S)PS19Vle/J; 1N4R tau) (The Jackson Laboratory, Cat #008169) were crossed into SORLA TG lines and SORLA KO lines; littermates from SORLA TG x PS19 and SORLA KO x PS19 were characterized at indicated timepoints. Mouse lines were housed with littermates with free access to food and water with a 12 hour light/day cycle. All procedures involving animals were performed under the guidelines of Sanford-Burnham Medical Research Institute (SBMRI) Institutional Animal Care and Use Committee.

### METHOD DETAILS

#### Immunoblot

For immunoblot analysis, 9MO WT, SORLA TG, PS19 and SORLA TG/PS19 mice were anesthetized and subjected to intracardial perfusion with 1xPBS. 7MO WT, SORLA KO, PS19 and SORLA KO/PS19 mice were anesthetized and subjected to intracardial perfusion with 1xPBS. Hippocampal tissues from each hemisphere were isolated, snap frozen on dry ice and stored at - 80°C. Lysates from hippocampal mouse tissue were extracted by homogenizer beads and resuspended in RIPA buffer in the presence of protease and phosphatase inhibitor cocktail. Protein concentration was measured using a Pierce™ BCA Protein Assay Kit. Proteins were separated by 4-20% Bio-Rad Criterion TGX Precast Gels, transferred onto PVDF membrances and blocked in 5% non-fat milk in 1xPBS. Blots were then probed with primary antibodies (mouse anti-SORLA, 1:1000; mouse anti-AT8, 1:1000; mouse anti-T13, 1:2000; mouse anti-HT7, 1:2000; mouse anti-Vinculin, 1:5000; Rabbit anti-APOE, 1:1000; Rabbit anti- β-Actin, 1:5000; rabbit anti-Iba1, 1:500; mouse anti-GFAP, 1:1000; goat anti-VPS35, 1:1000; rabbit anti-VPS26a, 1:1000; rabbit anti-VPS26b, 1:1000; rabbit anti-VPS29, 1:500) in 5% BSA/1xPBS overnight at 4°C, washed in 1xPBS with 0.1% tween-20, probed with HRP-conjugated secondary antibodies 2h at RT. After washing in 1xPBS, Blots were then incubated with ECL Chemiluminescent Substrate and immunoblot signals were acquired using a Chemidoc imaging system. For blot analysis, bands were quantified by Fiji ImageJ (NIH), targeted proteins were normalized to actin or vinculin and plotted relative to WT (set to

#### Immunohistochemistry

9MO WT, SORLA TG, PS19 and SORLA TG/PS19 mice were anesthetized and subjected to intracardial perfusion with 1xPBS. 9MO WT, SORLA KO, PS19 and SORLA KO/PS19 mice were anesthetized and subjected to intracardial perfusion with 1xPBS. Half hemi-brains were post-fixed with 4% PFA, dehydrated in 30% sucrose, embedded in O.C.T. and subjected to cryostat sectioning. Free-floating tissue sections (30µm) were rinsed in 1xPBS, incubated with blocking solution (0.5% Triton X-100, 5% BSA in PBS) for 2h at RT, immunostained with primary antibodies (mouse anti-AT8, 1:300; goat anti-Iba1, 1:200; rabbit anti-GFAP, 1:300; guinea pig anti-NeuN, 1:500; rat anti-CD68, 1:200; rabbit anti-C1q, 1:300; rat anti-Clec7a, 1:200; rabbit anti-P2y12, 1:200; guinea pig anti-PSD95, 1:300; rabbit anti-Vglut1, 1:400; rabbit anti-Plexin B1, 1:200; sheep anti-Plexin B2, 1:100, goat anti-GFAP, 1:300; rabbit anti-Iba1, 1:200) overnight at 4°C, then washed in PBS (3x) and subsequently incubated with Alexa Fluor Plus Secondary Antibodies (Invitrogen, 1:500) 2h at RT. After washing in 1xPBS, brain sections were mounted with DAPI Fluoromount-G and overlaid with glass coverslips. Images were acquired using a Zeiss confocal microscope (LSM 710) or APERIO ScanScope FL (Leica) system at 10x, 20x, or 60x magnification. For each comparison, all groups within a replicate experiment were stained simultaneously to avoid batch variation and imaged at the same fluorescent intensity. To quantify percent area covered, an optimal threshold was established for each stain in Fiji ImageJ and all the samples were quantified using matched threshold settings.

#### Volumetric analysis

Every tenth coronal brain section (30 um) starting from the appearance of hippocampus to the dorsal end of the hippocampus were mounted for each mouse. The brain slides were stained with 0.1% Sudan black in 70% ethanol at RT for 20 min, washed in 70% ethanol for 1 min (3x), washed in distilled water (3x) and mounted with Fluoromount-G and overlaid with a glass coverslip. Slides were scanned using an Aperio AT2 scnaner (Leica) at 10x magnification, ventricle and hippocampus area were measured using Aperio ImageScope Software (Leica).

#### Three-dimensional reconstructions

For microglia engulfment and synapse analysis, brain sections were imaged on a Zeiss LSM 880 microscope at 63x magnification with 0.39 µm z-sections. Image stacks with Iba1, PSD95 and Vglut1 or C1q-stained sections were used in three-dimensional reconstructions rendered using Imaris software 9.9. Vglut1 and PSD95 puncta within a size range of ∼0.5 to 1.1 µm were detected using the Imaris spot function to create discrete spheres with coordinates indicating center of mass; putative synapses were detected by quantification of pre- and postsynaptic spheres opposed to each other with a maximum distance of 1 μm (sum of the maximum puncta radius) according to published protocols (Fogarty, Hammond et al. 2013, Klenowski, Fogarty et al. 2015, Belmer, Klenowski et al. 2017). Iba1-positive microglia were 3D-reconstructed using the surface rendering function and vglut1, PSD-95, C1q puncta inside the microglia were quantified. 6-9 microglia within the hippocampal CA1 region were analyzed per mouse.

#### Sample preparation for proteomics analysis and LC-MS/MS

Proteomics analysis and LC-MS/MS was performed in the Sanford Burnham Prebys Bioinformatics Core. Frozen 9MO hippocampus tissue from WT, SORLA TG, PS19 and SORLA TG/PS19 mouse brain were homogenized and resuspended in RIPA buffer in the presence of protease and phosphatase inhibitor cocktail. Protein concentration was quantified using a Pierce™ BCA Protein Assay Kit. Proteins were then reduced by the addition of 5 mM Tris(2-carboxyethyl) phosphine (TCEP) at 30 °C for 60 min, followed by alkylation of cysteines with 15 mM iodoacetamide (IAA) for 30 minutes in the dark at room temperature. Urea concentration was reduced to 1 M by adding 50 mM ammonium bicarbonate. Samples were digested overnight with Lys-C/trypsin at room temperature with constant agitation at a 1:25 enzyme:protein ratio. Following digestion, samples were acidified using 0.1% FA and desalted using AssayMap C18 cartridges mounted on an Agilent AssayMap BRAVO liquid handling system. Cartridges were sequentially conditioned with 100% acetonitrile (ACN) and 0.1% FA; samples were then loaded, washed with 0.1% FA, and eluted with 60% ACN, 0.1% FA. Peptide concentration was determined using a NanoDrop spectrophotometer. Samples were subjected to mass spectrometry analysis using an EASY nanoLC system. Buffer A consisted of H2O/0.1% FA; Buffer B consisted of 80% ACN/0.1% FA. Samples were separated over a 90-min gradient of increasing Buffer B on analytical C18 Aurora column (75 μm × 250 mm, 1.6 μm particles; IonOpticks) at a flow rate of 300 nL/min. The mass spectrometer was operated in positive data-dependent acquisition mode, and the Thermo FAIMS Pro device was set to standard resolution with the temperature of FAIMS inner and outer electrodes set to 100 °C. A three-experiment method was set up where each experiment utilized a different FAIMS Pro compensation voltage: - 50, − 70, and − 80 Volts, and each of the three experiments had a 1 second cycle time. A high resolution MS1 scan in the Orbitrap (m/z range 350 to 1500, 60 k resolution at m/z 200, AGC 4e5 with maximum injection time of 50 ms, RF lens 30%) was collected in top speed mode with 1-second cycles for survey and MS/MS scans. For MS2 spectra, ions with charge states between + 2 and + 7 were isolated with quadruple mass filter using a 0.7 m/z isolation window, fragmented with higher-energy collisional dissociation (HCD) with normalized collision energy of 30% and the resulting fragments were detected in the ion trap as rapid scan mode with AGC of 5e4 and maximum injection time of 35 ms. The dynamic exclusion was set to 20 sec with a 10 ppm mass tolerance around the precursor.

#### Proteomic data analysis

Raw files were searched with SpectroMine software using default settings. The search criteria were set as follows: full tryptic specificity was identified where 2 missed cleavages were allowed, carbamidomethylation (C) was set as a fixed modification and oxidation (M) as a variable modification. The false identification rate was set to 1%. Spectra were searched against the curated Uniprot mus musculus database including human Tau protein sequence and common contaminants from the GPM cRAP sequences. Data was further processed using the MSstats package (version 4.2) in R (version 4.1.2). We avoided imputation of missing values prior to statistical testing using MSstats; instead we calculated pseudo Log2 fold-change (L2FC), adj.pvalue and pvalue of proteins completely missing in one condition in proteins with missing values. The imputed (pseudo) L2FC was calculated as the sum of intensities of the protein (i.e., sum of feature intensities of a given protein within a given sample) across all replicates of the same group that it was detected, divided by 3.3. For these proteins, the imputed pvalue and adj.pvalue was calculated by dividing 0.05 or 0.1, respectively, by the number of replicates that the given protein was confidently identified, and multiplied by the number of peptide features used for quantification in the given protein. Thus, the imputed L2FC provides an estimate of the protein abundance in the experimental conditions detected, while the imputed pvalue or adj.pvalue reports the confidence of the imputation and reflects the consistency of protein detection across replicates in the experimental group or conditions it was detected. Custom searches were performed for human SORLA (expressed in SORLA TG animals) and MAPT (human P301S tau).

Principal Component Analysis (PCA) was carried out in R version 4.1.2 with PCATools package (version 2.6.0) using log2 protein intensity for all significantly selected proteins summarized by dataProcess function from MSstats (version 4.2); multiple testing correction was performed with the Benjamini and Hochberg approach using MSstats. To calculate z-score values within each replicate, row (protein-wise) z-scores were computed in R version 4.0.2 using the scale function by subtracting mean intensity of each protein from the corresponding intensities of the biological replicates, and dividing the resulting values by the standard deviation of the intensities. GO (Gene Ontology) analysis of grouped DEP genelists was performed using GO DAVID (https://david.ncifcrf.gov/tools.jsp).

#### Electrophysiology

For analysis of long-term potentiation (LTP), ex-vivo hippocampal slices were prepared from 6-month-old WT, SORLA TG, PS19 and SORLA TG/PS19 mice as described previously (Huang, Zhao et al. 2017, Zhu, Liu et al. 2022). Under terminal isoflurane anesthesia, mice were transcardially perfused with ice-cold, sucrose-based artificial cerebrospinal fluid (aCSF) of the following composition: 190 mM sucrose, 25 mM D-glucose; 25 mM NaHCO3, 3 mM KCl, 1.25 mM NaH2PO4, 5 mM MgSO4, 10 NaCl mM, 0.5 mM CaCl2 saturated with carbogen (95% O2/5% CO2) and adjusted to pH 7.4 and 300 mOsm. After perfusion, mice were decapitated, and brains dissected and cut into a block of live brain tissue containing the hippocampus and cortex. A vibrating-blade microtome (Leica VT1200S) was used to cut 400 µm thick horizontal slices containing both cortex and hippocampus. Slices were transferred to a holding chamber containing warm (34°C) aCSF of the following composition: 125 mM NaCl, 25 mM NaHCO3, 3.0 mM KCl, 1.25 mM NaH2PO4, 2.0 mM CaCl2, 1.0 mM MgSO4, 10 mM D-glucose saturated with carbogen (95% O2/5% CO2) adjusted to pH 7.4 and 300 mOsm. Slices were left to recover at room temperature in oxygenated aCSF for 2 hours, transferred to a recording chamber and perfused with warm (32°C) oxygenated aCSF containing 100 µM picrotoxin (Palop, Chin et al. 2007) at a rate of 2 ml/min. For recording, both stimulating and recording electrodes were positioned within the dentate gyrus respectively on an upright microscope (Olympus). An electrical stimulation protocol was used to evoke field excitatory post synaptic potentials (fEPSP) in the middle of the molecular layer of the dentate gyrus. LTP was induced by two (2) trains of high frequency stimulation (HFS: 100 Hz for 1s at 20 s intervals) using a concentric bipolar stimulating electrode. fEPSPs were recorded using glass electrodes filled with aCSF and placed in the molecular layer of the dentate gyrus (dendritic region). All recordings of synaptic activity were carried out using a Multiclamp 700B and signals were filtered at 3 kHz, digitized and sampled using pClamp10 software. The magnitude of potentiation was calculated as percentage (%) change in fEPSP slope normalized to baseline values. The average change in synaptic potentiation at 1 h following induction of LTP was used to compare different experimental groups.

#### Nucleus isolation and snRNAseq library preparation

Three frozen 9MO hippocampus tissue from WT, PS19 and SORLA TG/PS19 groups were pooled and lysed in Miltenyi nuclei extraction buffer containing 0.2 U/µl RNase inhibitor using a gentleMACS Octo Dissociator (Program 4C_nuclei_1) according to the manufacturer protocol. Miltenyi Debris removal solution was applied to the nuclei suspension, and NeuN-AF488 and Olig2-AF647 were added to the samples and incubated on ice for 30 min. Sort buffer (1 U/ul RNase inhibitor, 1% BSA in PBS) was added to 4ml and DAPI was added at a 1:1000 dilution; the mixture was centrifuged at 1,500g for 5min and pelleted nuclei were re-suspended in 500 µl sort buffer. Nuclei were then vortexed before subjecting samples to FACS sorting. 150,000 nuclei were sorted into 90 µl collection buffer (5 U/µl RNase inhibitor, 10 % BSA in PBS). 10,000 nuclei were used to generate Gel Beads-in-emulsions and barcoded using the 10X Genomics Chromium Next GEM Single Cell 3′ Reagent Kit v3.1 (Dual Index) following manufacturer’s instructions. Libraries were sequenced to a median depth of approximately 50,000 reads per nucleus using an Illumina NovaSeq 6000 instrument at the UC San Diego IGM Genomics Center.

#### snRNAseq data processing, doublet detection and removal

Cell Ranger count was used to align samples to the reference genome (mm10), quantify reads, and filter reads with a quality score below 30. Aggregated and sequencing depth normalized count matrix from Cellranger aggregate *(cellranger aggr)* output was used for downstream analysis using Seurat version 4.0.5 (Hao, Hao et al. 2021) and R version 4.0.2. Lower quality cells with >10% mitochondrial reads and top 2% quantile of nFeature_RNA (number of detected genes) were removed. We used a two-step strategy to identify and remove potential doublets and multiplets. The Scrublet29 method over a random subset of 30,000 nuclei barcoded nuclei was used to identify clusters of doublets in the resulting snRNAseq library. Identification of heterotypic doublets comprising two different cell types was performed using Scrublet by simulating doublets, and through subsequent generation of nearest neighbor classifiers. During the first step, we applied the scrublet method version 0.2.3 (Wolock et al., 2019) to identify potential doublets for each sample, using the raw count matrix with default settings. During second step, we identified doublets/multiplets using scds package version 1.12.0 (Bais and Kostka 2020) in R version 4.2.1 as follows. The output of Cellranger aggregate was split into individual samples. Each sample was processed independently using Seurat pipeline to cluster cells. Briefly, raw data was normalized using *NormalizeData*. Top 2000 highly variable genes were identified using *FindVariableFeatures*. Data was scaled using *ScaleData*. Sequencing depth (nCount_RNA) and percent mitochondrial reads were regressed out during the *ScaleData* step. PCA on the scaled data was run using *RunPCA*. Cells were clustered using *RunUMAP*, *FindNeighbors*, and *FindClusters* with top 25 PCA dimensions and *resolution=0.5*. Seurat objects were converted to SingleCellExperiment objects using *as.SingleCellExperiment. cxds*, *bcds*, and *cxds_bcds_hybrid* functions were applied to the datasets to determine doublet scores. Cells with scds doublet score >0.75 were labeled as doublets and removed from downstream analysis. Data integration was performed using Seurat and Harmony version 0.1.0 (Korsunsky, Millard et al. 2019). Aggregated data with the doublets and lower quality cells removed was first normalized using *NormalizeData*. Top 2000 highly variable genes were identified using

*FindVariableFeatures*. Any mitochondrial or ribosomal genes and high expression genes (Malat1 and Xist) were removed from the list of highly variable genes. Data was scaled using *ScaleData* and the effects of sequencing depth (nCount_RNA) and mitochondrial percentage were regressed out. *RunPCA* was used to perform PCA on the scaled data and subsequently the samples were integrated using Harmony’s *RunHarmony* function with samples as the variable to remove. Cell clustering was performed using *RunUMAP*, *FindNeighbors*, and *FindClusters* with top 30 PCA dimensions and *resolution=0.2.* Cluster markers were determined using *FindAllMarkers* in WT sample only. Differential expression was performed using *FindMarkers* and MAST test. p-value correction for multiple testing using the Bonferroni method was performed using Seurat under default settings. GO (Gene Ontology) analysis of grouped DEG genelists was performed using GO DAVID (https://david.ncifcrf.gov/tools.jsp). To identify GO CC (Cellular Component) DEGs with little or no expression in microglia, transcripts (brainseq.org) that were expressed in less than 1% with an expression z-score <-1.33 in microglia were selected as transcripts of low abundance.

#### RNAscope analysis

Mice were anesthetized and perfused with cold 1xPBS, half-brain hemispheres were collected and stored in 10% neutral-buffered formalin 24h. Tissues were transferred to 70% ethanol and dehydrated in a graded series of ethanol and xylene, followed by infiltration by melted paraffin held at 60°C. FFPE Tissue blocks were cut into 5 μm sections on Fisher Scientific SuperFrost Plus Slides. Samples were prepared using the manufacturer’s instructions (RNAscope Multiplex Fluorescent Reagent Kit v2). After deparaffinizing, slides were incubated in hydrogen peroxide for 10 min at RT and washed in distilled water. Target retrieval was performed in a steamer for 20 min, and slides were subsequently washed in distilled water. For RNA-protein co-detection, diluted primary antibody (Goat anti-GFAP, 1:100; guinea pig Anti-NeuN, 1:100; rabbit anti-Iba1 Alexa 594, 1:50) was added to the slides and incubated overnight at 4°C; slides were washed in fresh PBST (1xPBS, 0.1% Tween 20) and fixed in 10% neutral-buffered formalin for 24h at RT. Slides were washed in fresh PBST, treated with Protease Plus at 40°C for 30 min, washed in distilled water, and incubated with the following RNAscope® Probes: Nrxn1, Cx3cr1, Slc17a7. mRNA probes were hybridized on slides in a HybEZ oven at 40°C for 2 hrs and washed with RNAscope wash buffer. Three additional hybridization steps with AMP1 (30 min), AMP2 (30 min), and AMP3 (15 min) solutions were applied to the slides with incubation in HybEZ at 40°C and rinsed with wash buffer between each step. Each individual probe signal was developed using TSA Vivid Dye. For a given channel (1, 2 and/or 3), HRP-C(n) was added to the slides (15 min, 40°C), followed by a diluted TSA Vivid Dye (520, 570 or 650; 30 min, 40°C), and HRP blocker (15 min, 40°C) with 2 buffer washes between each addition step. Fluorophore-conjugated secondary antibody diluted in co-detection antibody diluent (1:200) was applied to the slides for 30 min at RT, and slides were subsequently washed in fresh PBST. After incubation with DAPI, slides were mounted with ProLong Gold Antifade Mountant and fitted with a glass coverslip. All samples were stained concurrently and images were acquired using a Zeiss 880 Confocal Microscope at 63x magnification. For the number of mRNA dots, an optimal threshold was established for each stain in Fiji ImageJ and all the samples were quantified utilizing the same settings.

#### Soluble/insoluble tau extraction

Sarkosyl extraction was performed as previously described (Kim, Tadros et al. 2024). Cortex tissue was homogenized using a Dounce homogenizer in 10× (v/w) cold resuspension buffer (1× PBS containing 5 mM EDTA, protease and phosphatase inhibitor cocktail). Tissue debris was then removed by centrifugation at 2,000g for 5 min at 4 °C. For immunoblot analysis, a portion of the homogenate was diluted 1:1 with RIPA buffer (1× PBS containing 5 mM EDTA, protease and phosphatase inhibitor cocktails, 1% deoxycholate, 1% Triton X-100, and 0.5% SDS), and supernatant was collected after centrifugation at 15,500g for 20 min at 4 °C. For detergent-insoluble proteins, the remaining homogenate was mixed 1:1 with 2× RIPA buffer lacking SDS and centrifuged at 100,000g for 30 min at 4 °C. Supernatant (S1) was collected, and the pellet was resuspended in 1× RIPA buffer containing 1% Sarkosyl (without SDS) and incubated for 1 h at room temperature on an orbital shaker. The Sarkosyl-soluble fraction (S2) was obtained by centrifugation at 200,000g for 30 min at 4 °C. The final pellet was resuspended in 1× PBS by sonication (probe sonicator, 30% amplitude, 10 cycles of 2′ on/1′ off; P2). All fractions were stored at −80 °C until use for Western blotting.

#### Tau biosensor assay

Tau seeding was performed using a FRET biosensor HEK293T cell line stably expressing tau-RD P301S-CFP and P301S-YFP (tau-RD cells) as described previously (Parra Bravo, Krukowski et al. 2024). 10^5^ cells were plated on PDL-coated coverslips in a 24-well plate and cultured in DMEM media with 10% FBS and 1% Pen/Strep. The following day, the media was changed, and cells were seeded with brain lysate. Brain lysate was prepared by homogenizing tissue in 10 times the volume of PBS with 0.02% NaN3, 1% protease and phosphatase inhibitors cocktail. After homogenization, lysates were centrifuged at 21,000 g for 15 min at 4 °C. The supernatant was collected, and protein concentration was measured using the BCA assay. Cells were seeded with 7 μg of protein using Lipofectamine 3000 according to the manufacturer’s instructions. After 72 h of incubation, cells were washed and fixed in 4% PFA, washed and stained with DAPI, and imaged by confocal microscopy (Zeiss 880). Twelve images per condition from biological triplicate were taken for quantification using ImageJ.

#### Size Exclusion Chromatography (SEC)

Fresh whole mouse brain was homogenized using a mechanical homogenizer in 300µL of lysis buffer (20mM Tris HCl pH7.5, 5mM EDTA, 150mM NaCl and 1% DDM) containing a protease and phosphatase inhibitor cocktail and centrifuged at 28,000 g for 30 minutes at 4°C. Supernatants comprising 1 mL of enriched aggregates were injected onto a Superose 6 Increase 5/150 GL column (Cytiva) with a running buffer composed of 110mM NaCl, 20mM HEPES and 0.1% DDM and ran at 4°C. Absorbance was measured at 280 nm. Following sample application to the column, samples were eluted with 1.5x Column Volume (CV) running buffer. Fractions were collected after 0.2 CV at a fixed fractionation volume of 150µL, resulting in a total of twenty-five fractions collected. 13uls from each fraction was separated on 4-20% SDS-PAGE gradient gels for immunoblot analysis.

#### Primary neuron culture

Pups at postnatal day 0 (P0) were sacrificed for neuronal culture; forebrain was dissected in pre-cold Hibernate-A media (1% Pen/Strep) and was digested in 2.5% trypsin at 37 °C for 15 min, followed the addition of FBS and DNase. Digested tissue was filtered using 70 μm strainers and centrifuged at 300 g for 5 min. Cell pellets were resuspended in DMEM media with 10% FBS and 1% Pen/Strep, and seeded on poly-D-lysine (PDL)-coated plates for 3 hours, and subsequently changed to neuronal culture medium (neurobasal media with B-27 supplement, 1 × GlutaMAX supplement, and 1% Pen/Strep). Half of the media volume was changed every 3 days, and neurons at DIV14 were used for experiments. For tau uptake assay, recombinant 2N4R human tau 1–441 oligomerized in 30 μM heparin at 37 °C for 24 h. Tau oligomers were subsequently conjugated to Alex555 (“tau-555”) using an Alexa Fluor 555 Microscale Protein Labeling kit according to the manufacturer’s instructions. Tau oligomers were added to cells for 2 hours. For immunostaining, cells were fixed with 4% PFA for 20 min at room temperature after two washes with PBS. Cells were then permeabilized with 0.5% Triton X-100 in PBS for 5 minutes, blocked with 5% BSA in PBS for 1 hour, and incubated with primary antibodies (chicken anti-MAP2,1:1000; mouse anti- T13, 1:500; rabbit anti-EEA1,1:300; rabbit anti-RAB7,1:300; rabbit anti-LAMP1, 1:300) overnight at 4 °C. Cells were washed and incubated with Alexa Fluor-conjugated secondary antibodies for 1 hour. Cells were washed and stained with DAPI and imaged by confocal microscopy (Zeiss 880).

#### Primary microglia culture

Primary microglia culture was performed as previously described (Zhu, Liu et al. 2022). Pups at postnatal day 3 (P3) were used for microglia culture. Forebrain was dissected in pre-cold DMEM with 1% Pen/Strep and was treated with Papain at 37 °C for 30 min, followed by adding FBS and DNase. Digested tissue was filtered with 70 μm strainers and centrifuged at 300 × g for 5 min. The cell pellets were resuspended in DMEM media with 10% FBS and 1% Pen/Strep, cells were seeded in PDL-coated flasks. GM-CSF (25 ng/ml) was added into the cultures after 5 days and removed before harvesting. Microglial cells were harvested twice by shaking (200 rpm, 60 min) 10–14 days after plating and subjected to various treatments within 24 h of harvest.

#### Cell-cell interaction analysis

Cell-to-cell interactions (CCIs) were inferred by Cellphone DB (Efremova, Vento-Tormo et al. 2020) with the human CCI database (v5.0.0) mapped to mouse orthologs using biomaRt (Durinck, Spellman et al. 2009). Specifically, the “statistical_analysis_method” function was called using default parameters to infer CCIs with mean expression of interacting partners (ligand or receptor) being significantly enriched (p value <0.05), compared to random permutation. CCI inference was performed for each sample between pairwise cell type interactions comprising astrocytes, microglia and neurons. Significant CCIs between cell pairs with a p value <0.05 were used in visualization and selected as candidates for further validation.

#### GeoMx digital spatial profiling and analysis

Coronal sections from three 9MO SORLA KO, PS19 and SORLA KO/PS19 animals were mounted on slides. Tissue sections (5µm) were prepared by sequential deparaffinization, antigen retrieval (pH 9.0), 0.1μg/ml proteinase K digestion (15 min at 37°C), post fixation, followed by hybridization of tissue with ultraviolet photocleavable probes (Mouse Whole Transcriptome Atlas, NanoString Technologies, Inc) (overnight at 37°C) according to the manufacturer’s standard instructions. The tissue was then labeled with an anti-GFAP antibody conjugated to Alexa-532 (1:50 dilution), an Alexal-594-conjugated anti-Iba1 antibody (1: 50 dilution) and Syto13 (1:10 dilution). A separate slide was labeled with an anti-NeuN antibody conjugated to Alexa-647 (1:50 dilution) and Syto13 (1:10 dilution), and labeled slides were scanned on a GeoMx DSP instrument. Adjacent sections were stained to detect AT8 ptau. ROI selection was performed as follows: for GFAP or Iba1 ROI’s, geometric selections were drawn in the CA1 or CA3 region comprising the stratum pyramidale, stratum radiatum and stratum lacunosum, and within the granule cell layer (GCL)/hilus within the DG region. For NeuN ROI’s, tight geometric ROI selections were drawn within the CA1, CA3 or DG GCL. Utilizing the GeoMx segmentation tool, each ROI was then illuminated with a UV light to photocleave oligonucleotides probe barcodes from either GFAP+, Iba1+ or NeuN+ cells, which were then collected individually into single wells in a 96-well plate. Library preparation and sequencing were conducted at the SBP Genomics Core on an Element Biosciences AVITI sequencer. GeoMx raw sequencing data was processed using command-line GeoMx NGS Pipeline version 2.3.3.10. Processed data QC, filtering, and normalization was performed using NanoStringNCTools version 1.4.0, GeoMxTools version 3.0.1, and GeoMxWorkflows version 1.2.0 in R version 4.2.1(according to pipeline provided at https://bioconductor.org/packages/devel/workflows/vignettes/GeoMxWorkflows/inst/doc/GeomxTools_RNA-NGS_Analysis.html). Segment QC parameters were set as “minSegmentReads = 1000, percentTrimmed = 80, percentStitched = 80, percentAligned = 80, percentSaturation = 50, minNegativeCount = 1, maxNTCCount = 1000, minNuclei = 10, minArea = 1000”. Segments that failed the QC metrics were removed from downstream analysis. Segment gene detection limit was set at 5%, and data was normalized using the Q3 method. Normalized expression data for astrocytes (GFAP), neurons (NeuN), and microglia (Iba1) were batch corrected separately using removeBatchEffect from Limma version 3.52.4 (Ritchie, Phipson et al. 2015) using brain regions collected as batches. Differential expression analyses were performed using limFit and eBayes functions from Limma package.

### QUANTIFICATION AND STATISTICAL ANALYSIS

All statistical analysis in this study was performed using GraphPad Prism software and RStudio. Differences were assessed by paired or unpaired t tests, or one-way or two-way ANOVA where appropriate; a minimum of p<0.05 was statistically significant. All experiments in figures (including supplemental figures) were repeated at least three times (independent experiments) unless specified otherwise in the figure legends.

## Supplemental Figure legends

**Figure S1.**
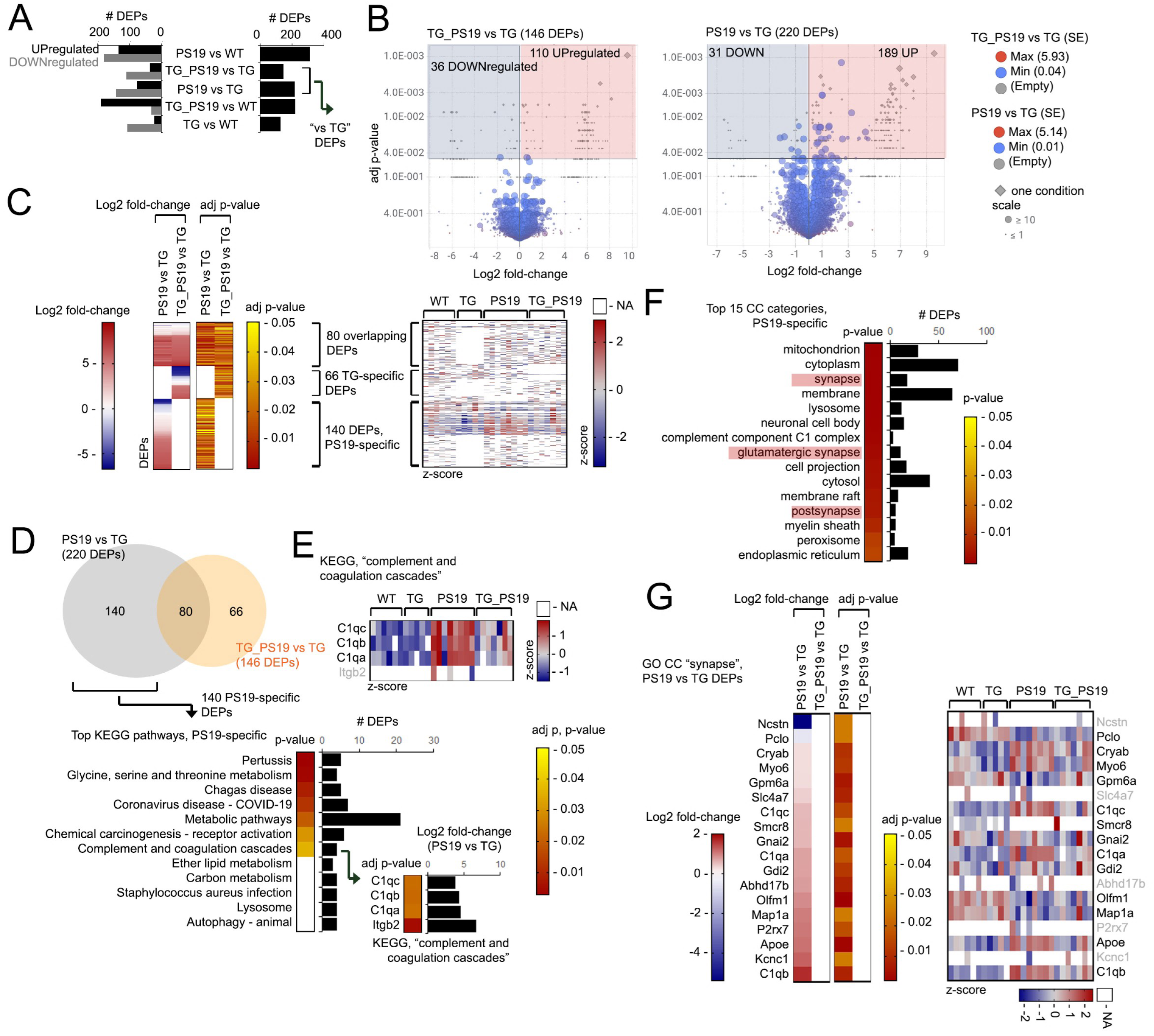
Characterizing effects of SORLA expression on PS19 mouse hippocampus. (A, B) (A) Total number (right graph) and number of upregulated (black bars) and downregulated (gray bars) DEPs (left graph), and (B) volcano plots depicting Log2 fold-change and adj p-values in “TG_PS19 vs TG” and “PS19 vs TG”, hippocampus. Number of DEPs (shown in brackets), as well as standard error (SE), frequency of detection are indicated by scaled circles; proteins absent under one condition are indicated by gray diamonds (legend) in (B). (C) Log2 fold-change and adj p-value in DEPs in “PS19 vs TG” and “TG_PS19 vs TG” comparisons indicated (left heatmaps), and z-score distribution for individual animals of the genotypes indicated (right heatmaps); “NA” values from missing peptide values are indicated in z-score heatmaps in white. (D) Venn diagram depicting overlapping DEGs between “PS19 vs TG” and “TG_PS19 vs TG” groups; p-value (heatmap) and number of DEPs (graph) in top 12 KEGG pathways by GO analysis of 140 PS19-specific is shown; adj-pvalue and Log2 fold-change (PS19 vs TG, graph) and adj p-value (heatmap) of genes in the KEGG “complement and coagulation cascades” pathway is shown in the inset. (E) z-score heatmap distribution of DEPs in the KEGG “complement and coagulation cascades” pathway from GO KEGG analysis in (D). (F) Top 15 GO CC categories in PS19- specific DEPs; p-value (heatmap) and number of DEPs (graphs) are shown. Categories related to “synapse” and synaptic function are highlighted. (G) Log2 fold-change and adj p-value (left heatmaps) of DEP comparisons indicated and z-score distribution of GO CC “synapse” DEPs in (F) in hippocampus from genotypes indicated (right heatmap). A high number of missing values in replicate animals (indicated in white/NA, light gray text labels) were observed in Itgb2 (E) and Ncstn, Slc4a7, Abhd17b, P2rx7 and Kcnc1 (G), indicating a low confidence level for these DEPs due to inconsistent detection amongst replicates. Related to Figure 2.

**Figure S2.**
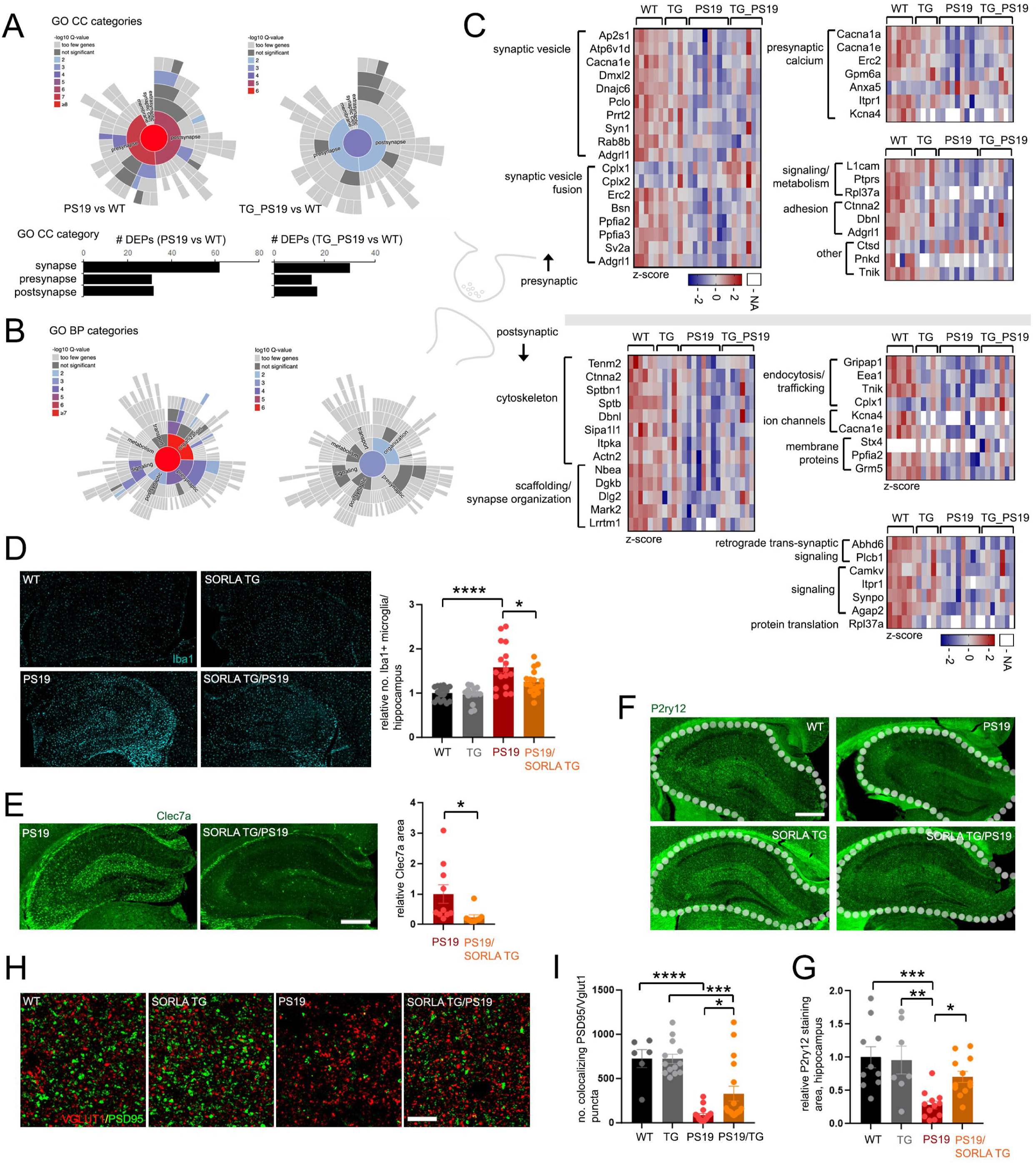
Characterizing synaptic and microglial changes in SORLA TG/PS19 vs PS19 hippocampus. (AC) Changes in synaptic components in PS19 and SORLA TG/PS19 hippocampus identified by proteomic analysis. Distribution of synaptic DEPs in PS19 vs WT, and TG_PS19 (SORLA TG/PS19) vs WT comparisons in GO cellular component (CC) (A) or biological process (BP) (B) synapse-related categories, identified using the synGO analysis tool (syngoportal.org). Colored heat maps indicate significance (-Log10 q-value) for DEPs in various GO CC and BP synapse-related categories shown, and the number of total synapse, pre-synaptic or postsynaptic DEPs in PS19 vs WT or TG_PS19 vs WT comparisons are indicated in bar graphs shown in (A). (C) Zscore distribution of presynaptic (upper panels) and postsynaptic (lower panels) DEPs from WT, TG (SORLA TG), PS19, and TG_PS19 hippocampus annotated by synGO analysis. Presynaptic and postsynaptic DEPs are clustered into various GO categories indicated. (D) Representative images of 9MO WT, SORLA TG, PS19 or SORLA TG/PS19 hippocampus stained for Iba1 (cyan). Adjacent graph shows quantified relative number of Iba1- stained microglia per hippocampus (normalized to WT, set to 1.0) (WT: 9F, 9M; STG: 8F, 6M; PS19: 10F, 7M; STG/PS19: 10F, 7M). (E) Representative image of 9MO PS19 and SORLA TG/PS19 hippocampus stained for Clec7a (green), bar = 500um. Adjacent graph depicts % Clec7a staining area in hippocampus (% Clec7a staining area/total hippocampus area) (PS19: 6F, 4M; STG/PS19: 5F, 4M). (F, G) (F) Representative images of P2ry12 (green) staining in 9MO WT, PS19, SORLA TG or SORLA TG/PS19 hippocampus, bar = 500um. (G) P2ry12 staining area in hippocampus quantified from images in (F) within the hippocampal area indicated by white border markings (WT: 5F, 5M; STG: 5F, 2M; PS19: 8F, 4M; STG/PS19: 8F, 5M). (H, I) Hippocampal sections from 9MO animals as indicated stained for Vglut1 (red)/PSD95 (green) and visualized for co-localizing Vglut1/PSD95 puncta, bar=10um. Adjacent graph depicts the number of co-localizing Vglut1/PSD95 puncta quantified using IMARIS (WT: 3F, 3M; STG: 8F, 5M; PS19: 10F, 7M; STG/PS19: 10F, 7M). All graphs represent mean±SE, where individual plots represent values from one animal. Statistical significance was determined by unpaired Student’s t-test (E) or One-way ANOVA with Tukey’s multiple comparisons (D), (G) and (I), *p<0.05, **p<0.01, ***p<0.001, ****p<0.0001. Related to Figure 3.

**Figure S3.**
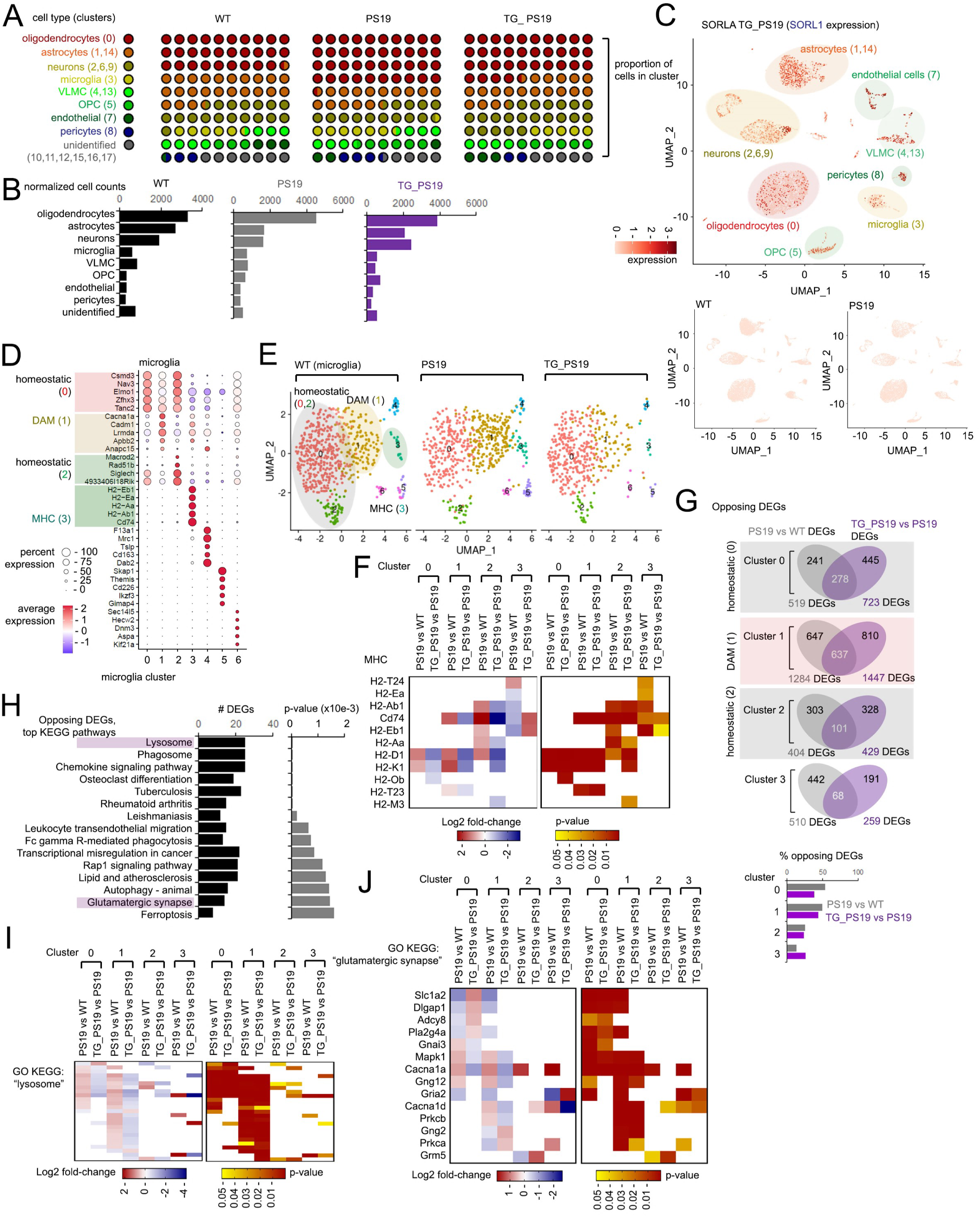
**Characterization of microglia snRNAseq profiles from PS19 and SORLA TG/PS19 hippocampus**. (A, B) 10x10 plot (A) and bar graph (B) showing relative distribution of cell types indicated in nuclei identified from 9MO WT, PS19 and SORLA TG/PS19 hippocampus. (C) Umap comprising all clustered nuclei from CNS cell types indicated in WT, PS19 and TG_PS19 hippocampus; heat-scales indicate expression of the human SORL1 transgene (“expression”). No human SORL1 expression was detected in WT and PS19 mouse hippocampus. (D) Top 5 microglia genes defining the clusters indicated (clusters 0 to 6). Homeostatic (clusters 0 and 2), DAM (cluster 1) and MHC (cluster 3) subclusters are indicated. (E) Umap feature plots depicting microglia from WT, PS19 and TG_PS19 hippocampus; colored indicator labels mark homeostatic (red, green), DAM (dark yellow) and MHC (light green) subclusters. (F) Heatmaps depicting Log2 fold-change (left heatmap) and p-value (right heatmap) of MHC-related DEGs in reclustered microglia nuclei from Clusters 0 through 3. (G) Venn diagram depicting opposing DEG expression profiles in microglia Clusters 0 through 3 in PS19 (PS19 vs WT) and SORLA TG/PS19 (TG_PS19 vs PS19) hippocampus. Bar graph shows percentage of DEGs reversed in PS19 (PS19 vs WT, gray) and TG_PS19 (TG_PS19 vs PS19, purple) microglia in the clusters indicated. (H) Top 15 GO KEGG pathways observed in PS19 microglia DEGs showing opposing expression profiles in SORLA TG/PS19 hippocampus. Bar graphs depict the number of DEGs and p-values for each KEGG pathway identified. (I, J) Expression profiles in opposing microglial DEGs associated with “lysosome” and “glutamatergic synapse” GO KEGG pathways. Heatmaps depict Log2 fold-change (left heatmaps) and p-value (right heatmaps) for lysosome (I) and glutamatergic synapse (J) DEGs in microglia Clusters 0 through 3. Related to Figure 4.

**Figure S4.**
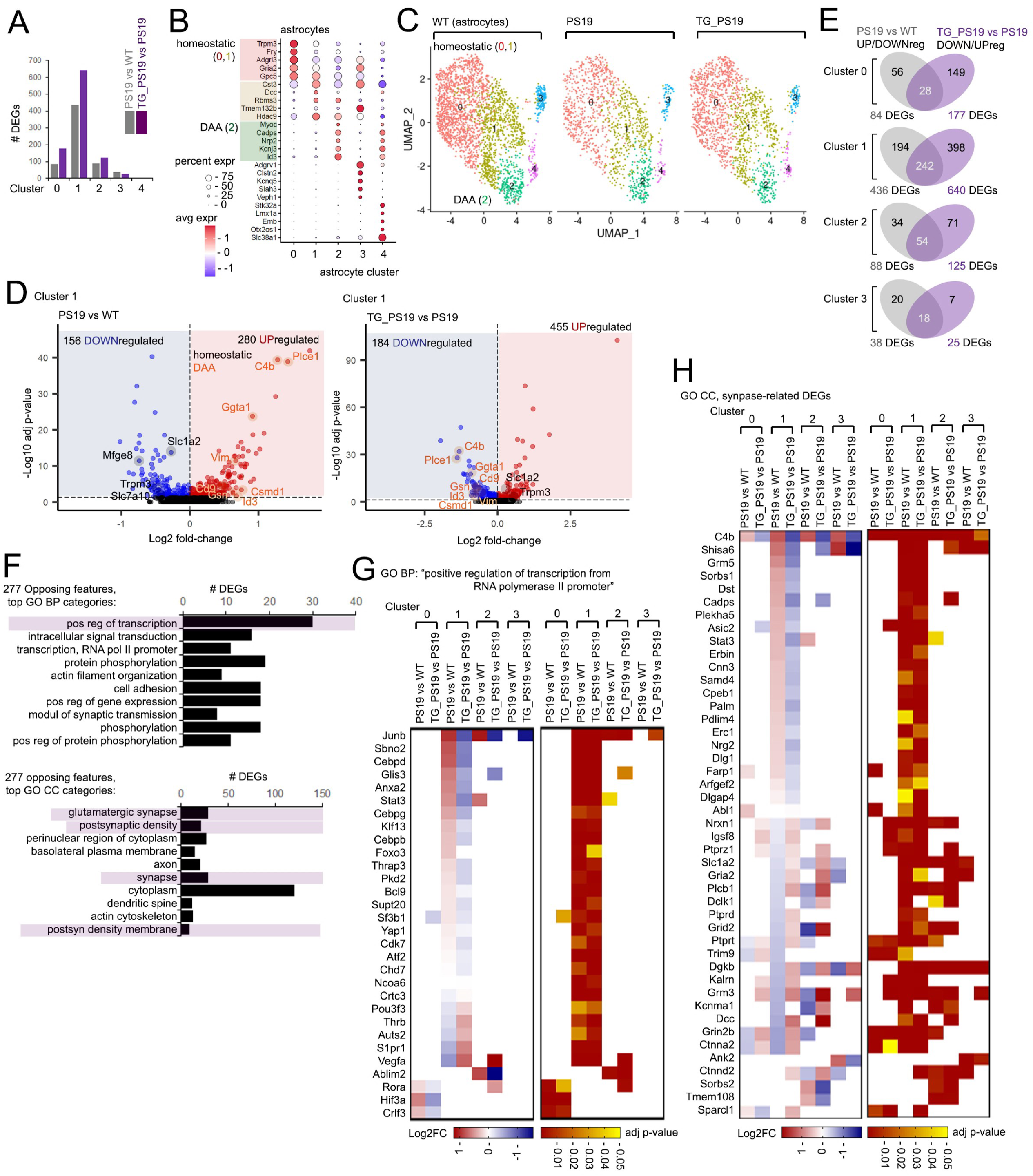
**SORLA upregulation reverses transcriptomic profiles in PS19 astrocytes**. (A) Number of DEGs identified from PS19 vs WT (gray) and TG_PS19 vs PS19 (purple) comparisons in astrocyte Clusters 0 through 4. (B) Top 5 astrocyte genes defining the clusters indicated (clusters 0 to 4). Homeostatic (clusters 0,1) DAA (cluster 2) subclusters are indicated. (C) Umap feature plots depicting microglia from WT, PS19 and TG_PS19 hippocampus; homeostatic (0,1 – red, yellow) and DAA (2 – green) subclusters are indicated. (D) Volcano plots depicting up and downregulated DEGs in PS19 vs WT and TG_PS19 vs PS19 comparisons in astrocyte Cluster 1 (adj p-value<0.05). Homeostatic (black) and DAM (orange) DEGs featuring opposing expression profiles in PS19 and TG_PS19 microglia are indicated. (E) Venn diagram showing opposing DEGs (“opposing features”) in PS19 astrocytes (PS19 vs WT) reversed in SORLA TG hippocampus (TG_PS19 vs PS19) in Clusters 0 through 3. (F) Top 10 GO BP (top graphs) and GO CC (bottom graphs) categories observed in 277 opposing expression features identified in (E). Number of DEGs (black bars) and p-values (gray bars) are shown in the bar graphs. (G, H) Expression profiles in opposing astrocyte DEGs associated with GO BP “positive regulation of transcription from RNA polymerase II promoter” (G) and GO CC synapse-related DEGs (H). Heatmaps depict Log2 foldchange (left heatmaps) and adj p-value (right heatmaps) in astrocyte Clusters 0 through 3. Related to Figure 5.

**Figure S5.**
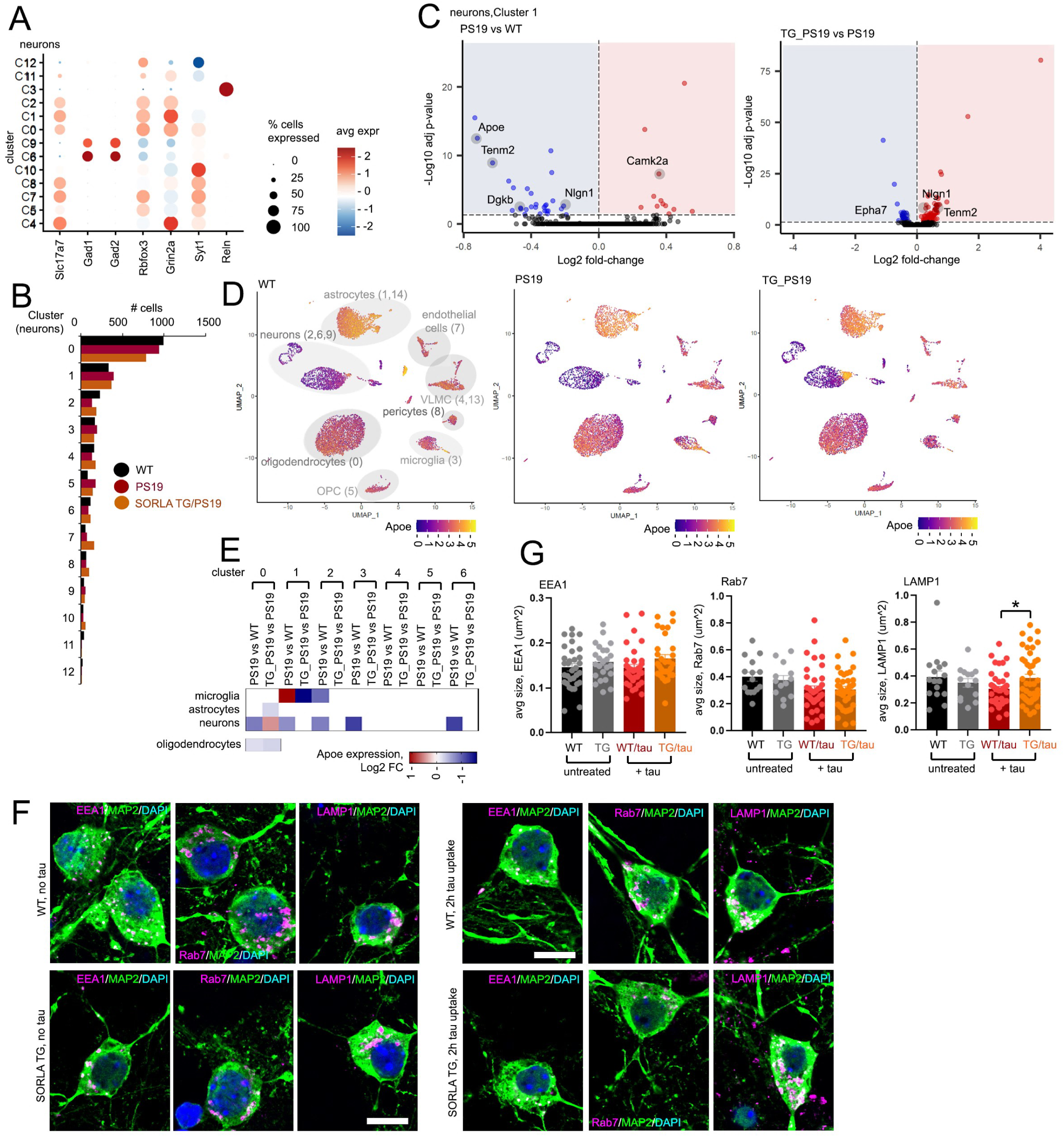
Reversal of PS19 expression profiles with SORLA upregulation in neurons. (A) Dot plot featuring cluster-specific expression markers in neurons; scales for percentage cell expression (plot size) and expression (red/blue heatmap scale) are shown on the right. (B) Bar graph showing number of cells observed in WT, PS19 and TG_PS19 neurons from each neuronal subcluster. (C) Volcano plots for DEGs from PS19 vs WT, and TG_PS19 vs PS19 comparisons in neuron Cluster 1 (adjp<0.05); DEGs related to “glutamatergic synapse” are indicated in gray. (D, E) Apoe expression in various CNS cell types. (D) Heat-scaled umap feature plot depicting Apoe expression in WT, PS19 and TG_PS19 hippocampus by snRNA-seq analysis. Annotated cell types are indicated in the WT umap feature plot. (E) Heatmap summary showing cluster-specific changes in Apoe expression in neurons and glia in PS19 vs WT, and TG_PS19 vs PS19 comparisons. (F, G) Characterizing effects of tau oligomers on cultured neurons. (F) Representative images of untreated DIV14 WT or SORLA TG neurons (left panels), or neurons treated 2h with tau oligomers (right panels) and stained for endolysosomal markers (EEA1, Rab7, LAMP1; purple), MAP2 (green) or nuclei (DAPI, blue) as indicated, bar=10um. (G) Quantification of EEA1, Rab7 or LAMP1 size in WT and SORLA TG (TG) neurons under untreated or tau-treated conditions. Graphs in (G) represent mean±SE, statistical significance was determined by One-way ANOVA, *p<0.05. Related to Figure 6.

**Figure S6.**
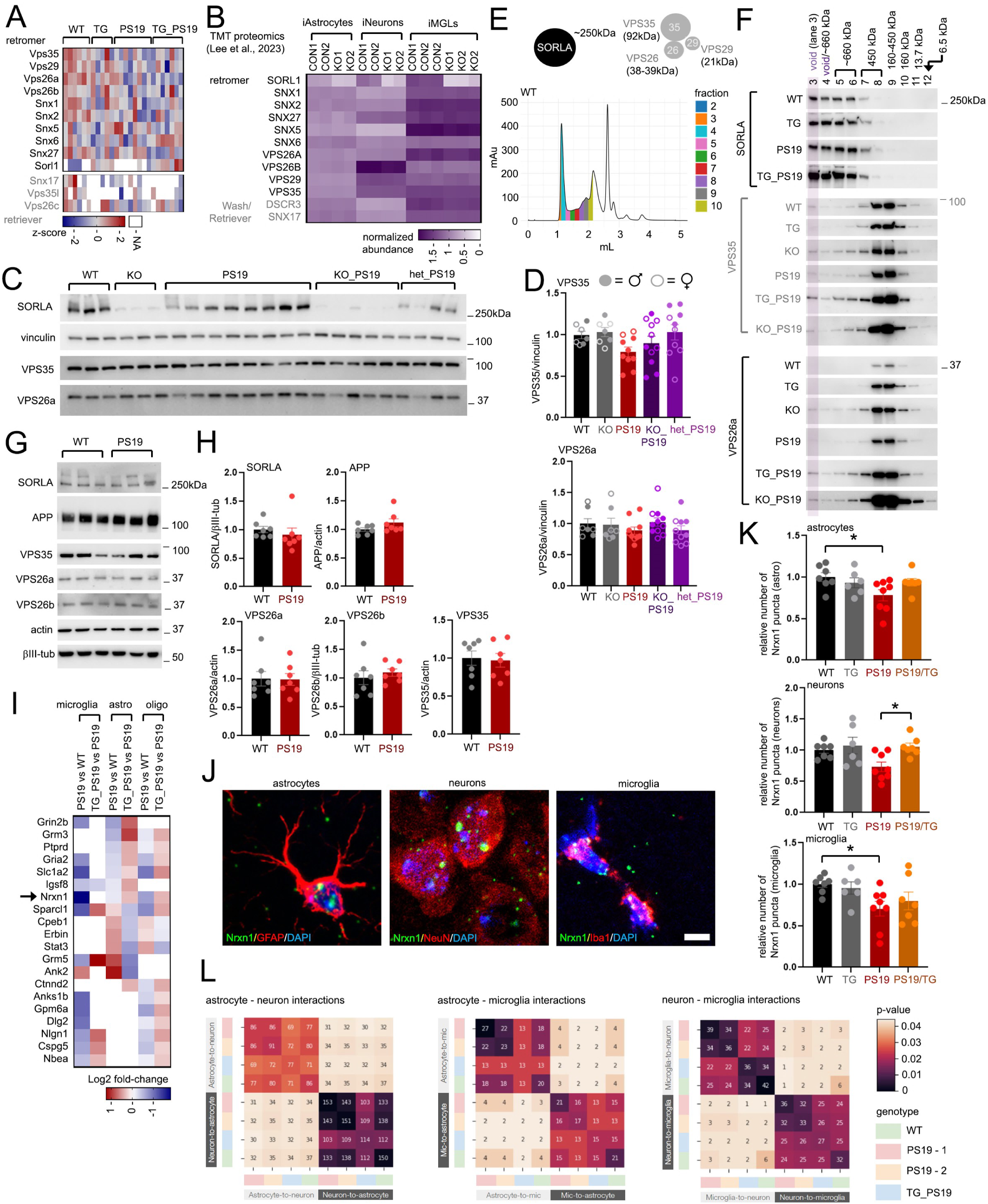
Effects of SORLA modulation on the retromer complex and synaptic targets. (A) Heatmap depicting z-score profiles of retromer/retriever components identified in 9MO WT, SORLA TG (TG), PS19, and SORLA TG/PS19 (TG_PS19) hippocampus. (B) Relative abundance of retromer and retriever components identified in iPSC-derived control and SORLA KO astrocytes, neurons and iMGLs as characterized by TMT-proteomics (Lee et al., 2023). (C, D) Characterization of high molecular weight SORLA and retromer complexes in mouse brain. (C) Schematic (upper panel), predicted molecular weight of SORLA and core retromer components. Lower panel showing SEC elution profiles of WT mouse brain; fractions are color-coded in the elution fractions, milli-Absorbance units (mAu) within the elution profiles are shown. (D) Elution profiles of SORLA, VPS35 and VPS25a in 1.5MO WT, TG, KO, PS19, SORLA TG/PS19 (TG_PS19) and SORLA KO/PS19 (homozygous KO_PS19) mouse brain. Void volume (fraction 3 and a portion of fraction 4) is indicated. (E, F) (E) Immunoblot analysis of retromer levels in 7MO mouse WT, KO, PS19, KO_PS19 and SORLA het KO/PS19 (het_PS19) hippocampus. (F) Quantification of VPS35 (upper graph) and VPS26a (lower graph) levels in mouse hippocampus from (E). Male and female mice represent individual plots as indicated. (G, H) (G) Representative immunoblot of retromer components in DIV14 cultured neurons. (H) Quantification of retromer components and APP from immunoblot in (G). (I) Heatmap (Log2 fold-change) showing expression profiles in synapse-related DEGs in microglia, astrocytes and oligodendrocytes. (J) RNAscope analysis of Nrxn1 expression in 9 month-old WT, SORLA TG (TG), PS19 and SORLA TG/PS19 (PS19/TG) hippocampus. Representative images of GFAP (red), Nrxn1 mRNA (green) in astrocytes (left panel), NeuN (red), Nrxn1 mRNA (green) in neurons (middle panel) and Iba1 (red), Nrxn1 mRNA (green) in microglia (right panel). Scale bar, 5um. (K) Graphs depict mean and SE of relative number of Nrxn1 puncta in stained cell types in (J) normalized to WT (set to 1.0). Individual plots represent average number of puncta quantified from one animal. Statistical significance was determined by Oneway ANOVA with Sidak’s multiple comparison, *p<0.05. (L) Number of shared interactions identified between different cell types between WT (green), PS19 (red and orange each mouse) and TG_PS19 (blue) animals. Related to Figure 7.

**Figure S7.**
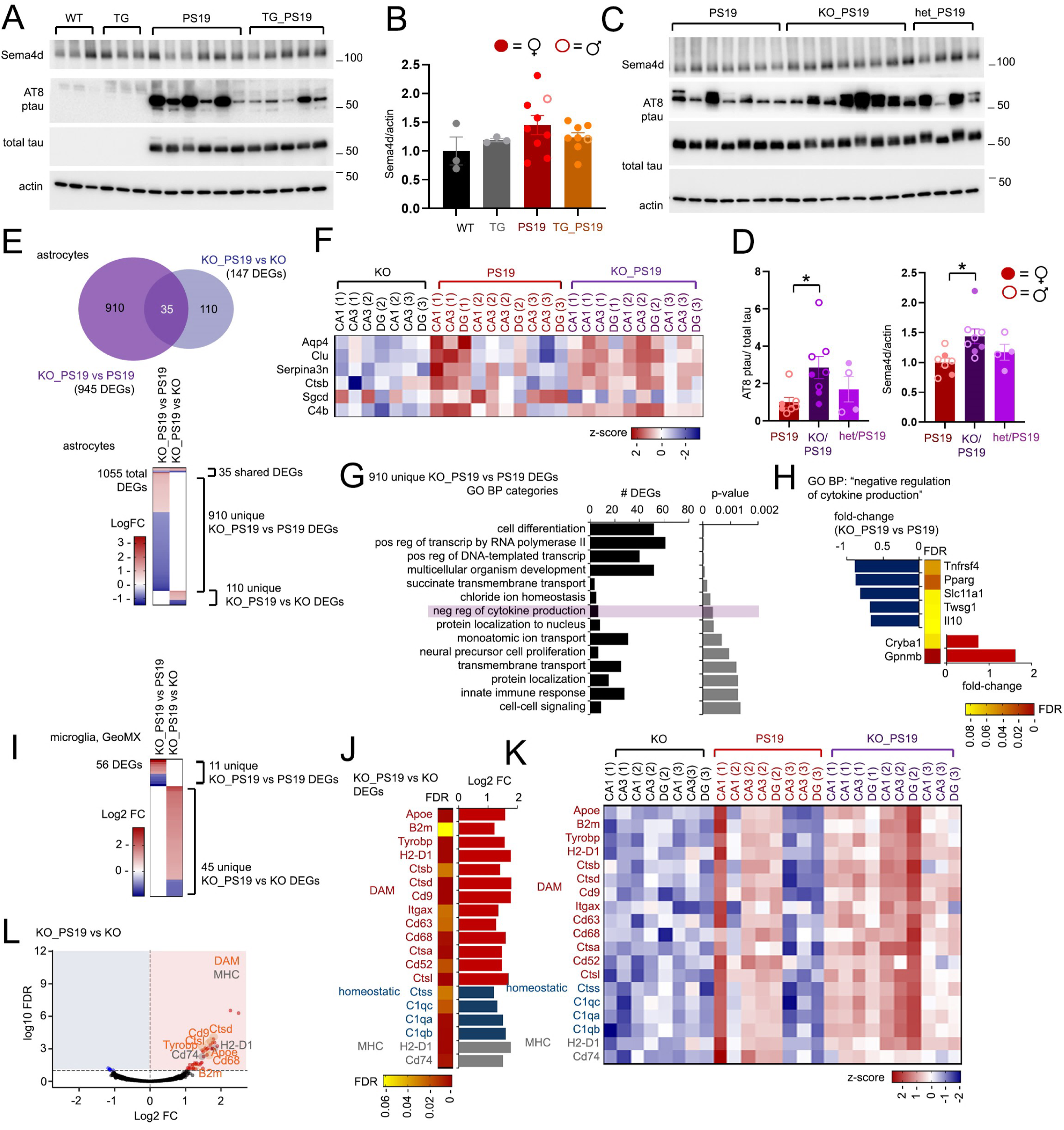
**Characterizing effects of SORLA modulation on the Sema4d/PlxnB pathway in PS19 mouse hippocampus**. (A) Immunoblots from 9MO WT, SORLA TG, PS19 or SORLA TG/PS19 (TG_PS19) hippocampal protein lysates to detect Sema4D, AT8 ptau, total human tau, or actin. (B) Quantification of Sema4D band intensity from blots in (A) normalized over actin, compared to WT (set to 1.0). (C) Immunoblots from 7MO WT, homozygous SORLA KO (“KO”), PS19, homozygous SORLA KO/PS19 (“KO_PS19”) or heterozygous SORLA/PS19 (“het_PS19”) hippocampal protein lysates to detect Sema4D, AT8 ptau, total human tau, or actin. (D) Quantification of AT8 ptau/total tau (left graph) or Sem4d band intensity normalized over actin (right graph) from blots in (C), compared to PS19 (set to 1.0). (E-H) Comparative transcriptomic analysis in SORLA KO/PS19 astrocytes. (E) Venn diagram (top) and heatmap (bottom) showing total number of unique and overlapping DEGs identified in SORLA KO/PS19 (“KO_PS19”) vs PS19 comparisons, or KO_PS19 vs SORLA KO (“KO”) comparisons. (F) Heatmap depicting z-scores for DAA DEGs in astrocytes from CA1, CA3 and DG ROIs in KO, PS19 or KO_PS19 hippocampus. ROIs from one of three individual animals per genotype are indicated in brackets. (G) GO analysis of 910 unique KO_PS19 vs PS19 DEGs; top 14 GO BP categories are shown. (H) Bar graphs depicting Log2 fold-change in KO_PS19 vs PS19 GO BP “negative regulation of cytokine production” DEGs. Scaled heatmap depicts FDR for DEGs identified. (I-L) Characterizing changes in microglia in KO_PS19 brain by GeoMX analysis. (I) Heatmap depicting DEGs from KO_PS19 vs PS19 or KO_PS19 vs KO comparisons in microglia. (J) Log2 fold-change (bar graph) and FDR (heatmap) of DAM, homeostatic and MHC-associated DEGs in microglia. (K) Heatmap showing z-score expression profiles of DAM, homeostatic and MHC genes in hippocampus from the genotypes indicated. Hippocampal subregions, and replicate animals (in brackets) are indicated. (L) Volcano plots from microglia KO_PS19 vs KO DEGs; representative DAM (orange) and MHC (gray) DEGs are indicated. Quantification in graphs from (B) and (D), data represent mean±SE and individual plots represent quantification from an individual animal. Statistical significance was determined by One-way ANOVA with Tukey’s multiple comparison (*p<0.05). Related to Figure 8.

**Figure S8.**
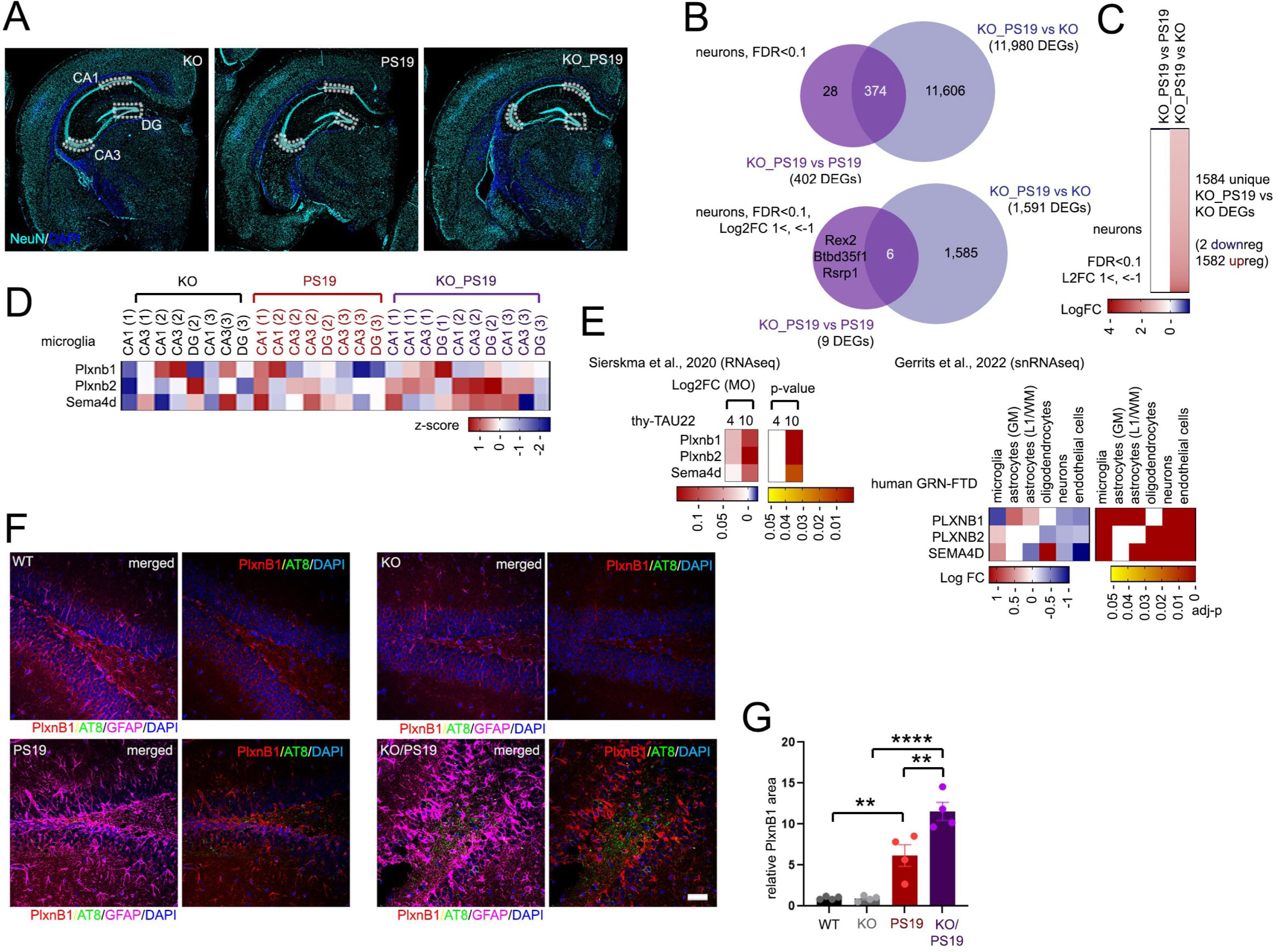
SORLA deletion exacerbates astrocyte and microglia activation in PS19 hippocampus. (A) Brain hemisections stained for NeuN (turquoise) and nuclei (DAPI, blue); ROI selections within CA1, CA3 and DG regions in 9MO KO, PS19 and KO_PS19 hippocampus for GeoMX analysis are indicated. (B) Venn diagrams showing overlap in KO_PS19 vs PS19 and KO_PS19 vs KO DEGs in neurons at FDR<0.1 (top) and FDR<0.1, Log2 fold-change >1, <-1 (bottom) cutoffs. (C) Heatmap of neuronal DEGs (FDR <0.1, Log2 fold-change >1, <- 1) in KO_PS19 vs PS19 and KO_PS19 vs KO DEGs. Unique KO_PS19 vs KO DEGs are almost exclusively upregulated (1582 DEGs), with only 2 DEGs downregulated. (D) Heatmaps depicting z-score expression profiles for Plxnb1, Plxnb2 and Sema4d in microglia in sampled hippocampal regions in KO, PS19 and KO_PS19 animals. Replicate animals are indicated in brackets. (E) Changes in PlxnB1/B2 and Sema4D expression observed in tauopathy and neurodegeneration. Left heatmaps: *Plxnb1/Plxnb2* and *Sema4d* expression levels in 4 or 10 monthold thy-TAU22 mouse brain (vs control) by bulk RNAseq analysis (Sierskma et al., 2020). Heat-scaled changes are indicated for Log2 fold-change (left), or p-value (right heatmap). Right heatmaps: PLXNB1, PLXNB2, or SEMA4D expression in human GRN-FTD brain vs control by snRNAseq analysis (Gerrits et al., 2022). Heat-scaled changes are indicated for Log fold-change (left), or adjusted p-value (right heatmap). (F) Representative DG images in WT, KO, PS19 or KO/PS19 hippocampus stained for PlxnB1 (red), AT8 ptau (green), and nuclei (blue, DAPI) (bar=50um). (G) Quantification of PlxnB1in DG images from (F) (WT: 1F, 3M; STG: 1F, 3M; PS19: 2F, 2M; SKO/PS19: 1F, 3M). Data represent mean±SE and individual plots represent quantification from an individual animal. Statistical significance was determined by One-way ANOVA with Tukey’s multiple comparison in (G), (**p<0.01, ****p<0.0001).

## Supplemental Tables S1-S4

Table S1, related to Figure 2.

Label-free proteomic analysis, 9MO WT, SORLA TG, PS19, SORLA TG/PS19 hippocampus

Sheet 1 - "L2FC_significance", DEPs (adjp<0.05)

Sheet 2 - z-score values across individual animals for significant DEPs

Table S2, related to Figure 2.

GO analysis, proteomics

Sheet 1 - "GO_CC_227 PS19 DEPs", GO CC (cellular component) categories, 227 DEPs unique to PS19 vs WT comparisons

Sheet 2 - "KEGG_140 PS19vsTG DEPs", GO KEGG pathways from 140 DEPs specific to PS19 vs TG comparisons

Sheet 3 - "GO_CC_140 PS19vsTG DEPs", GO CC categories from 140 DEPs specific to PS19 vs TG comparisons

Table S3, related to Figures 4-6.

snRNA-seq analysis, 9MO WT, PS19 and SORLA TG/PS19 hippocampus Sheet 1 - Microglia, clusters C0 to C3, L2FC>0.3,<-0.3; p<0.05

Sheet 2 - Astrocytes, clusters C0 to C4, adjp<0.05 Sheet 3 - Neurons, clusters C0 to C12, adjp<0.05 Sheet 4 - Oligodendrocyte DEGs, adjp<0.05

Table S4, related to Figure 8.

GeoMx analysis (DEGs, z-scores), 9MO SORLA KO, KO/PS19, PS19 hippocampus DEGs and z-scores for astrocytes, microglia and neurons are shown

## Notes

### Competing Interest Statement

The authors have declared no competing interest.

